# Patterns of variation in canal and root number in human post-canine teeth

**DOI:** 10.1101/2022.02.01.478656

**Authors:** Jason J. Gellis, Robert A. Foley

## Abstract

Descriptive morphology of tooth roots traditionally focuses on number of canals and roots. However, how or if canal and root number are related is poorly understood. While it is often assumed that canal number is concomitant with root number and morphology, in practice canal number and morphology do not always covary with external root features. To investigate the relationship between canal and root number, fully developed, adult post-canine teeth were examined and quantified from medical computerized tomography scans from a global sample of 945 modern humans. We tested the hypotheses that root and canal number do not follow a 1:1 ratio, that canal to root ratios differ between teeth, and that canal to root ratios differ across populations. Results indicate that not only is root number dependent on canal number, but that this relationship become more variable as canal number increases, varies both between individual teeth and by population, and changes as populations increase in distance from Sub-Saharan Africa. These results show that the ratio of canal number to root number is an important indicator of variation in dental phenotypes.

## Introduction

Tooth root anatomy varies in canal and root number, and canal number does not always covary with root number. Various aspects of this have been studied in modern humans (Kovacs, 1971; Ackerman et al., 1973; Vertucci and Gegauff, 1979; Hsu and Kim, 1997; Zorbaa et al., 2014; Ahmed et al., 2017), extant hominoids (Kupczik et al., 2005; Emonet et al., 2012; Moore et al., 2013, 2015), and fossil hominins (Wood et al., 1988a; Plavcan and Daegling, 2006; Kupczik et al., 2009; Kupczik and Hublin, 2010; Le Cabec et al., 2013; Moore et al., 2016). However, the numerical relationship between canals and roots is poorly understood. This study uses medical CT scans to investigate the relationship and variability between canal and root number of fully developed, adult post-canine teeth in a global sample of modern humans (n = 945 individuals) from several archaeological/osteological collections. Specifically, we asked (1) what is the relationship between root number and canal number; (2) does this relationship vary by tooth; and (3) does the relationship between canal and root number vary in global groups?

### Root and canal formation

Tooth canal and root formation are comprised of a series of reciprocal cellular interactions in the dental papilla of the developing tooth (Jernvall and Thesleff, 2000). Central to the process, is Hertwig’s epithelial root sheath (HERS), which is derived from the cervical loop of the enamel organ and is thought to be responsible for root number, shape and length (Nanci, 2012; Luder, 2015). Following crown formation, mesenchyme cells form the blood vessels, nerves, and connective tissue of the pulp cavity and root canals (Wright, 2007). Simultaneously, the HERS extends apically, interacting with the mesenchyme cells of the developing canal structures, and differentiating into odontoblasts responsible for dentin and cementum production (Li et al., 2017).

During root morphogenesis, the HERS produces inter-radicular processes (IRP’s), finger-like protrusions adjacent the cervical foramen of the tooth crown. The extension and fusion of opposing IRPs across the cervical foramen create multiple secondary foramina which, in turn, form multiple tooth roots (Kovacs, 1971; Orban and Bhaskar, 1980); and it may be that number and orientation of IRP’s are responsible for the variation in canal and root forms (Figure 1). While molecular regulation and tooth morphogenesis have been extensively studied in tooth crowns, the mechanisms responsible for variation in canal and root structures is poorly understood. Because of its extensive role in root formation, HERS has been an area of focus; and several studies have shown that disturbances in formation of the HERS results in abnormalities in root number and shape (see Luder, 2015 for a review).

**Figure 1:**
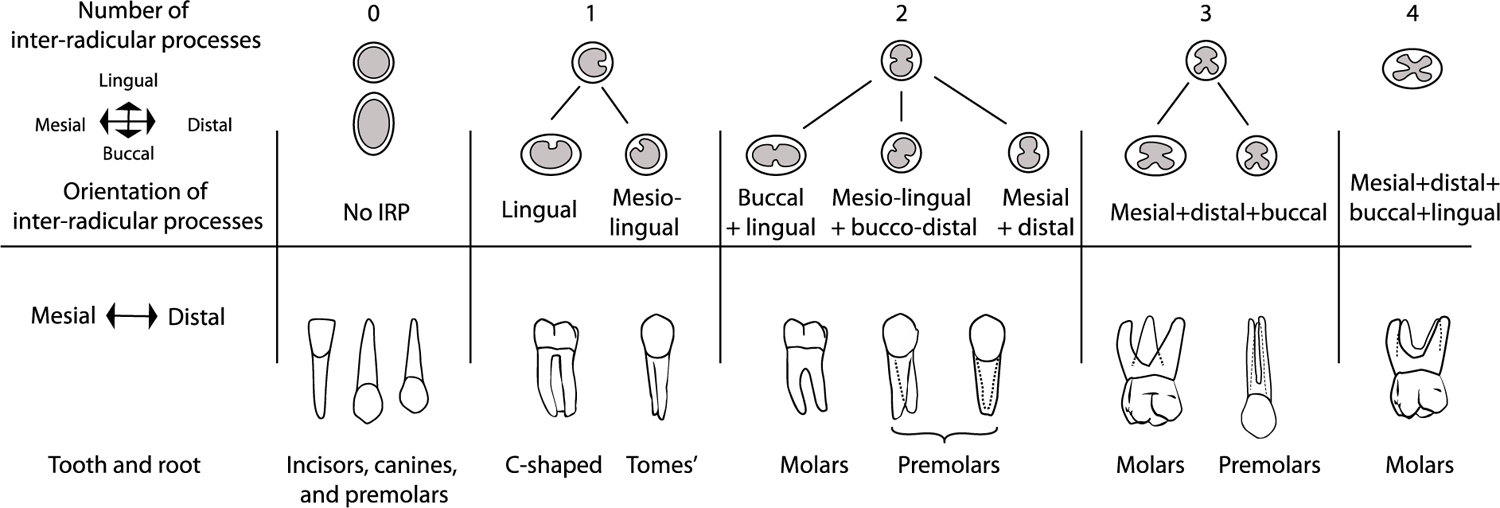
**Top:** The location and on the apical foramen of the tooth crown where the inter-radicular processes (IRP’s) form determines the number and orientation of each tooth root/tooth roots. For example, in a tooth with mesial and distal roots, two inter-radicular processes arise from the buccal and lingual borders of the apical foramen, forming mesial and distal secondary apical foramina upon fusion. Grey = apical foramina of the developing tooth crown. **Bottom:** Fully developed roots of different types of teeth with the same number, but different orientation of IRPs. From right to left: single rooted teeth, single rooted teeth in which IRP did not fuse with opposing side of apical foramen, 2 rooted teeth in which 2 opposing IRPs fused, 3 rooted teeth in which 3 opposing IRPs fused, 4 rooted teeth in which 4 opposing IRPs fused.

Though morphogenesis of internal and external root structures are concurrent processes, the completed structures do not always covary. There is great variation and complexity in root canals. While it is easy to conceptualize canals as round holes which taper towards the roots’ apex, in reality, many teeth have multiple canals of differing shape and orientation within a single root. These canals can join and separate in unpredictable places and the more ovoid the cross-section the greater the propensity for complexity (Vertucci and Gegauff, 1979; De Pablo et al., 2010; Ahmed et al., 2017). Possible causes of divergence in canal and root number have been attributed to uneven deposition of dentin on the walls of the canal (Manning, 1990), trauma to the HERS by radiation or chemical interference (Fischischweiger and Clausnitzer, 1988), and/or failure of the HERS to fuse on different sides of the root (Nelson and Ash, 2010; Nanci, 2012).

In this paper we (1) test the hypothesis that there is no difference between canal and root number in the pooled post-canine teeth in our sample; (2) test the relationship between canal and root number in the individual post-canine teeth of the jaws; (3) test the relationship between canal and root number in pooled and individual teeth, by geographical regions.

## Materials and Methods

### Dental formula

Categorically premolars with P, and molars with M. Tooth numbers are labelled with super- and subscripts to differentiate the teeth of the maxilla and mandible respectively. For example, M^1^ indicates the 1st maxillary molar while M_1_ indicates the 1st mandibular molar.. Through the course of evolution, apes and old world monkeys have lost the first and second premolars of their evolutionary ancestors (Novacek, 1986; White et al., 2012), thus the remaining 2 premolars are numbered 3 and 4.

### Human samples

The 945 individuals used in this study were recovered from archaeological sites across the globe (Figure 2). These individuals are stored in osteological collections at the Smithsonian National Museum of Natural History, Washington D.C., USA (SI), American Museum of Natural History, New York, USA (AMNH), and the Duckworth Laboratory (DW) at the University of Cambridge, England (summarized in Figure 3). Only adult individuals, based on the eruption, occlusion, and closed root apices of M^3^/M_3_’s (or M^2^/M_2_’s in the case of congenitally absent M^3^/M_3_’s), were used in this study.

**Figure 2:**
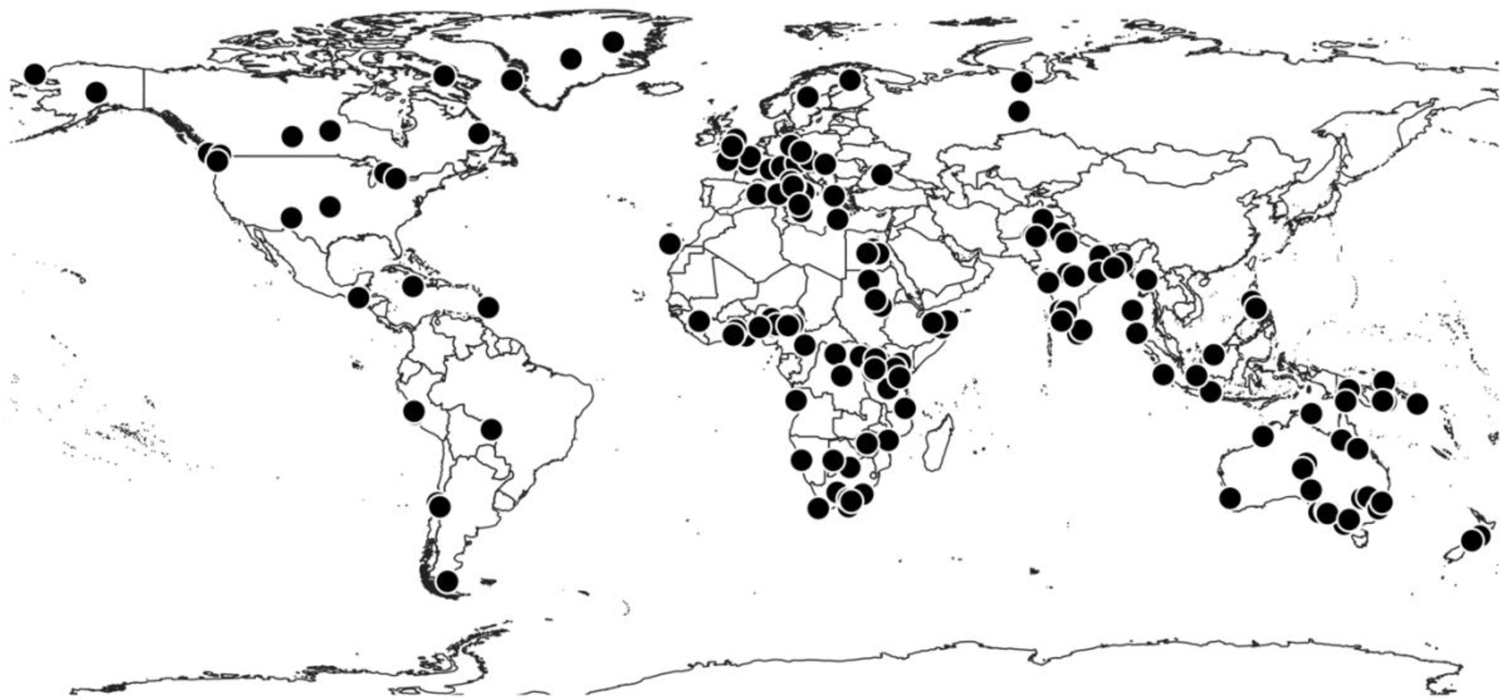
Map of archaeological sites for individuals used in this study.

**Figure 3:**
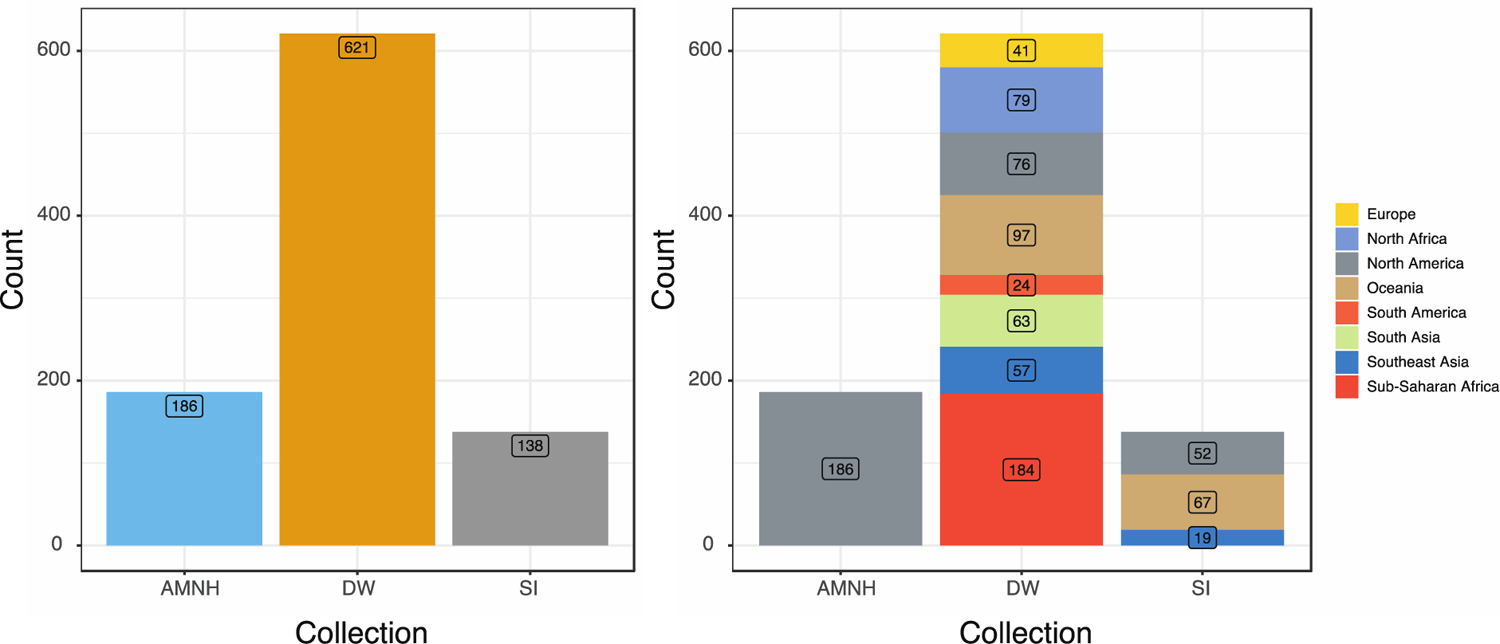
Human population sample sizes by collection. **Left:** Bar plot of counts for entire sample (n = 945). **Right:** Counts of samples divided up by collection, and geographic locations given by collection records. A complete list of the individuals used in this study, their collection information, antiquity, sex, and locality based on available records is listed in Supplementary Materials Table A.

### American Museum of Natural History

The 186 individuals from the AMNH collection are comprised of humans from Point Hope, Alaska, North America (Figure 3, right). These individuals, are attributed to the Ipiutak (500 BCE – 500 CE) and Tigara (1300-1700 CE) cultures (Rainey, 1941, 1947, 1971; Larsen and Rainey, 1948). Information on sex (Figure 4) and antiquity come from the AMNH archives and publications associated with the collection (ibid).

**Figure 4:**
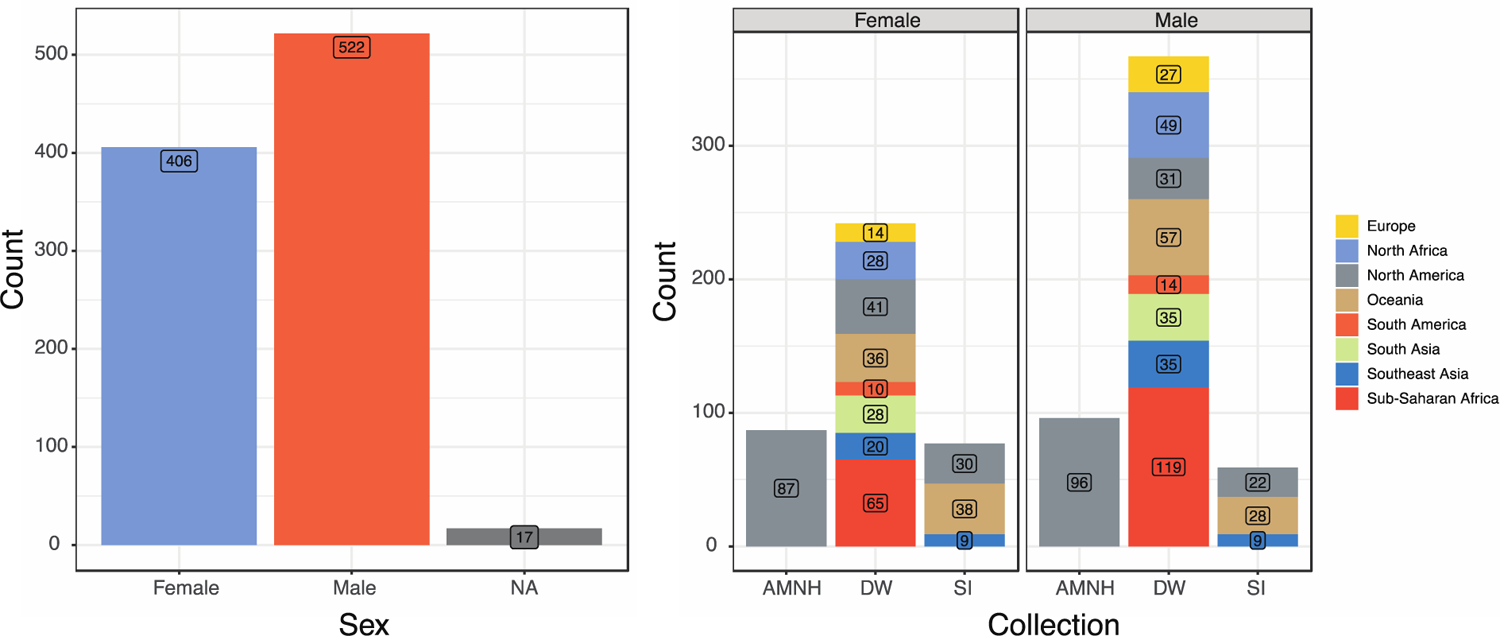
Human population sample sizes by location and sex. **Left:** Bar plot of sex for entire sample (n = 945). **Right:** Sex divided up by collection and geographic locations given by collection records. Individuals of undetermined sex (‘NA’) are not included in the plot on the right to improve readability. They are: AMNH (NA = 3), DW (NA = 12), and SI (NA = 2). A complete list of the individuals used in this study, their collection information, antiquity, sex, and locality based on available records is Supplementary Materials Table A.

### Duckworth Laboratory

The majority of individuals (n = 621) used in this study come from the DW Laboratory collections (Figure 3, right). The DW is comprised of several private collections as well as research collections from the University of Cambridge Departments of Zoology, Anatomy, and Museum of Archaeology and Anatomy (Mirazón-Lahr, 2011). The oldest individuals used in this study come from the archaeological sites of Badari, Egypt (4000-3200 BCE), Jebel Moya, Sudan (100 BCE-500 CE) and Ngada, Egypt (4400-4000 BCE), in North-East Africa. The majority of the remaining individuals are ∼200 years old. In many cases information on exact locality, age, and age of death, is unavailable. Information on sex (Figure 4) comes from DW archives. A complete list of the DW individuals used in this study, their collection information, antiquity, sex, and locality based on available records is listed in Supplementary Materials Table A.

### Smithsonian National Museum of Natural History

The 138 individuals from the SI collection are from Oceania, Southeast Asia, and Greenland. Individuals from Oceania belong (n = 67) to four different populations: Australia (Aboriginal), New Zealand (Maori), the Philippines, and Papua New Guinea (Figure 3, right). Individuals from Southeast Asia (n = 19) are from Indonesia. Inuit individuals comes from the North-West coast of Greenland (n = 52). While all SI individuals were recovered from archaeological sites, information on exact locality, age, and age of death, is unavailable. However, information on sex taken from archives at the SI (Figure 4) as reported in Copes (2012).

The populations in this study have been, at their broadest level, grouped into five major human populations: Sub-Saharan Africa, West-Eurasia, Sahul-Pacific, Sunda-Pacific, and Sino-Americas (Table 1 & Figure 5). Though Table 1 reports information for sex, this is for descriptive purposes. All analyses and reported results are for pooled sex samples. A complete list and description of individuals included in this study is listed in the Supplementary Materials Table A.

**Figure 5:**
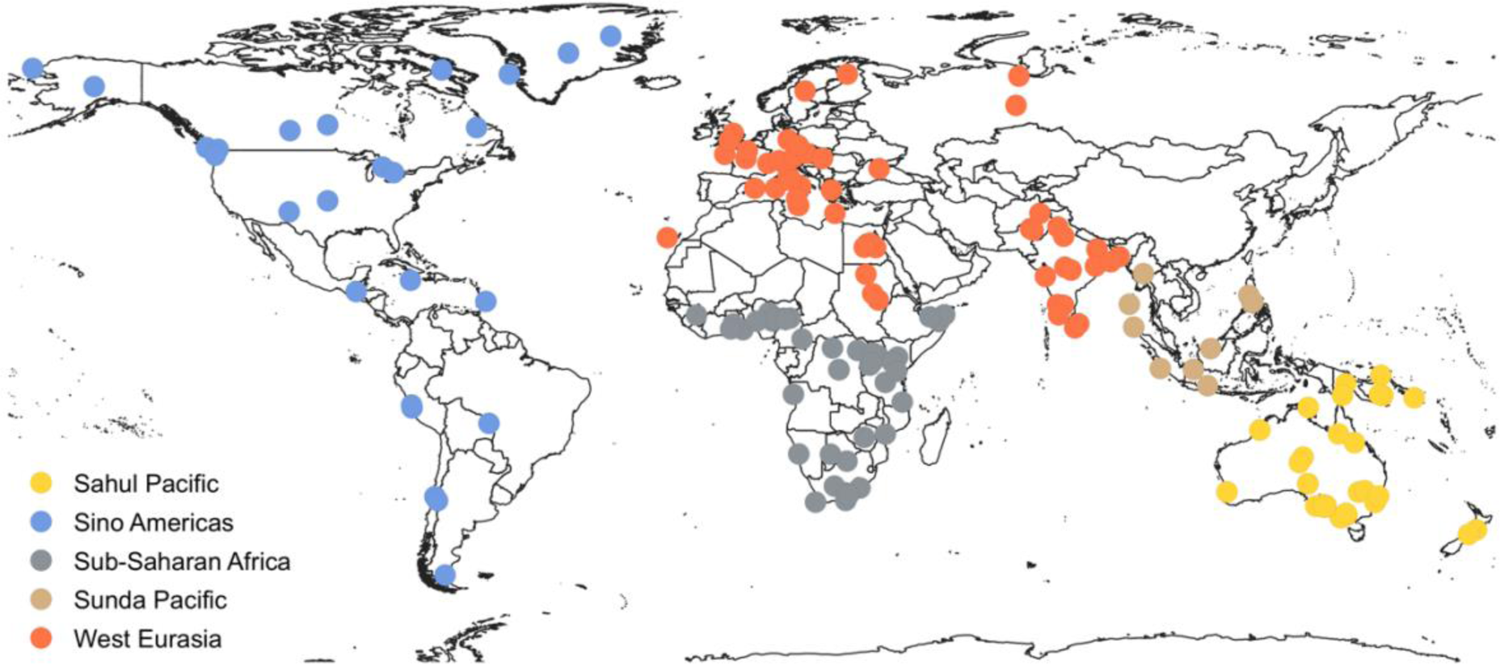
Map of archaeological sites for individuals used in this dissertation adapted to show the five major human populations.

**Table 1:** Sample populations used in this study

| Populations* | Male | Female | Unknown | Total |
| --- | --- | --- | --- | --- |
| Sahul-Pacific | 84 | 74 | 9 | 167 |
| Sunda-Pacific | 42 | 28 | 3 | 73 |
| Sub-Saharan Africa | 119 | 65 | - | 184 |
| West Eurasia | 111 | 70 | 2 | 183 |
| Sino-Americas | 163 | 168 | 7 | 338 |
| Total | 519 | 405 | 21 | 945 |

These groupings are derived from two major works. The first is Cavalli-Sforza’s *The History and Geography of Human Genes* (1994), a synthesis of global genetics with nearly half a century’s worth of geographical, ecological, linguistic, archaeological, and paleoanthropological research. Among the author’s many conclusions are that all available evidence points to 1) an African origin for *H. sapiens; and* 2) the fact that a series of dispersal and admixture events can classify and map where major human populations (as listed above) and their subsequent subdivisions originated and dispersed through the ancient world. The volume (1994:317) also recognises that dental data “on northern Asia, southeast Asia, and the Americas are generally in excellent agreement with those from single genes.” The dental data they refer to are crown and root trait frequencies collected and analysed by Christy Turner (Nichol et al., 1984; Turner II, 1986, 1987, 1989). These data, along with later core collected works on dental crown traits and biogeography utilising the ASUDAS (Scott, 1988; Turner II et al., 1991; Stringer et al., 1997; Irish, 1998; Hanihara, 2013; Scott et al., 2018), form the second basis for major human geographical subdivisions presented here. These researchers (ibid) have shown that teeth are effective for identifying the same prehistoric population identities and movements discussed by Cavalli-Sforza (1994), as well as capturing the phenotypic diversity within populations, and the differences that arise between them after extended periods of isolation. The most current collections of dental anthropological research (Rathmann et al., 2017; Scott et al., 2018; Rathmann and Reyes-Centeno, 2020) are increasingly in accordance with the most recent genomic studies (Pickrell and Reich, 2014; Fu et al., 2016; Skoglund et al., 2016; Rathmann et al., 2017; Posth et al., 2018; Reich, 2018), further reinforcing the utility of teeth as phenotypic records of human biogeography and evolutionary history.

### Use of cone beam computed tomography for visualizing internal and external features of tooth roots

In clinical settings (e.g., dental, hospital, etc.), cone beam computed tomography (CBCT or CT) is widely utilized to visualize internal and external structures of the crown and root(s) (see Martins and Versiani (2018) for an in-depth discussion of this topic). An important parameter supporting reliability of visualization for the study of root and canal anatomy is voxel size. In 3D medical imaging a single voxel is a cubic representation of a single value of space within a cubic volume. For example, a hypothetical 300×300×300 cubic volume would have 27,000,000 voxels. Thus, the lower the voxel size relative to the volume of 3D CBCT, the greater the resolution. Compared to micro-CT (µCT) which operates on the micron scale (a thousandth of a millimetre) for increased resolution, CBCT uses larger voxel sizes at the millimetre scale which results in a relatively decreased resolution. However, CBCT has been proven to be reliable for detecting root an canal number and morphology in specific teeth or individual roots (Blattner et al., 2010; Michetti et al., 2010; Domark et al., 2013; Pécora et al., 2013). Maret et al. (2014) compared in vitro CBCT images of different voxel sizes (76, 200, and 300 μm) with µCT (41 µm) and observed discrepancies of hard tissue morphology (i.e. cervical margins, cusp tips, incisal edges) were only significant at 300µm (P = .01, Wilcoxon test). These studies (additionally, see Martins and Versiani (2018) for an extensive overview of CBCT and µCT on root canal anatomy by tooth) have shown that CBCT can clearly and accurately detect structures such as root number, canal number, and configuration of the main root canal systems.

### Imaging of osteological collections

Following the method developed by Gellis and Foley (2021) we used cone-beam computed tomography (CT) to analyse 5,970 post-canine teeth (Table 2) from the right sides of the maxillary and mandibular dental arcades of individuals (n= 945) from a global sample of humans (Table 1). While information for all teeth from both sides of the maxillary and mandibular arcades was recorded, only the right sides were analysed to avoid issues with asymmetry and artificially inflated sample size. Full skulls of specimens from the SI and AMNH were scanned by Dr. Lynn Copes (Copes, 2012) using a Siemens Somatom spiral scanner (70 µA, 110 kV, slice thickness 1.0 mm, reconstruction 0.5 mm, voxel size mm^3: 0.5×0.5×0.3676). Full skulls from the DC were scanned by Professor Marta Miraźon-Lahr and Dr. Frances Rivera (Rivera and Mirazón-Lahr, 2017) using a Siemens Somatom Definition Flash scanner at Addenbrooke’s Hospital, Cambridge England (80µA, 120kV, slice thickness 0.6mm, voxel size mm^3: 0.3906×0.3906×0.3). For all collections, crania and mandibles were oriented on the rotation stage, with the coronal plane orthogonal to the x-ray source and detector. Permission to use the scans has been granted by Dr. Copes, Professor Miraźon-Lahr and Dr. Rivera.

**Table 2:**
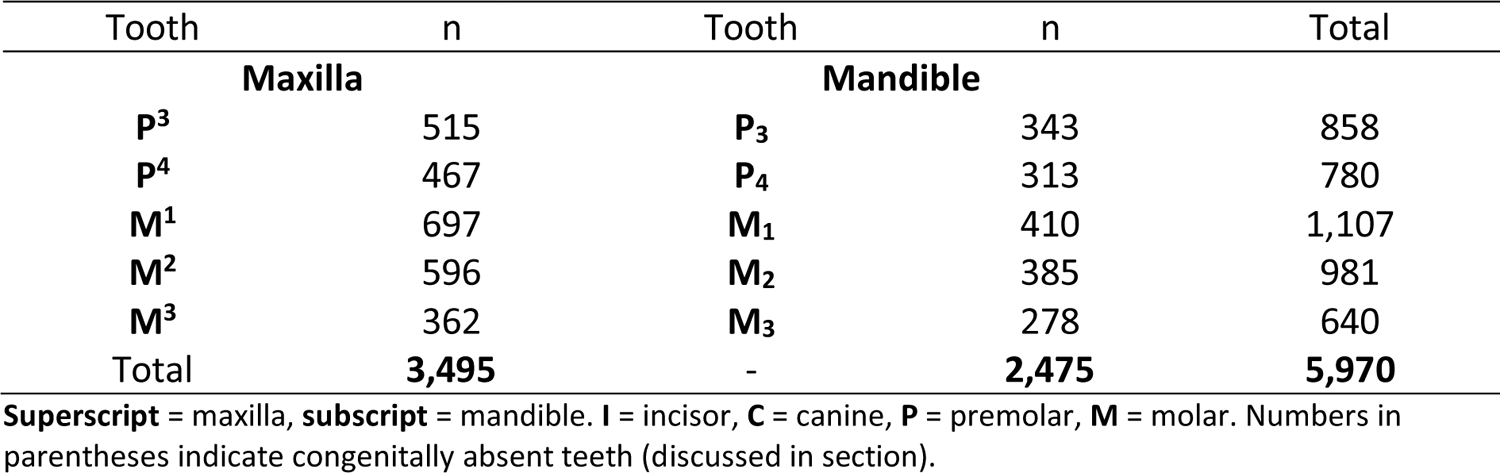
Tooth counts of the right side of the maxillary and mandibular dental arcades.

| Tooth | n | Tooth | n | Total |
| --- | --- | --- | --- | --- |
| Maxilla |  | Mandible |  |  |
| **P<sup>3</sup>** | 515 | **P<sub>3</sub>** | 343 | 858 |
| **P<sup>4</sup>** | 467 | **P<sub>4</sub>** | 313 | 780 |
| **M<sup>1</sup>** | 697 | **M<sub>1</sub>** | 410 | 1,107 |
| **M<sup>2</sup>** | 596 | **M<sub>2</sub>** | 385 | 981 |
| **M<sup>3</sup>** | 362 | **M<sub>3</sub>** | 278 | 640 |
| Total | **3,495** | - | **2,475** | **5,970** |

**Superscript** = maxilla, **subscript** = mandible. **I** = incisor, **C** = canine, **P** = premolar, **M** = molar. Numbers in parentheses indicate congenitally absent teeth (discussed in section).

### Analysis of CT images

Transverse CT cross sections of roots and canals were assessed in the coronal, axial, and sagittal planes across the CT stack, using measurement tools in the Horos Project Dicom Viewer (Figure 6) version 3.5.5 (https://www.horosproject.org, 2016). Only permanent teeth with completely developed roots and closed root apices were used for this study. While information for all teeth from both sides of the maxillary and mandibular arcades was recorded, only the right sides were analysed to avoid issues with asymmetry and artificially inflated sample size.

**Figure 6:**
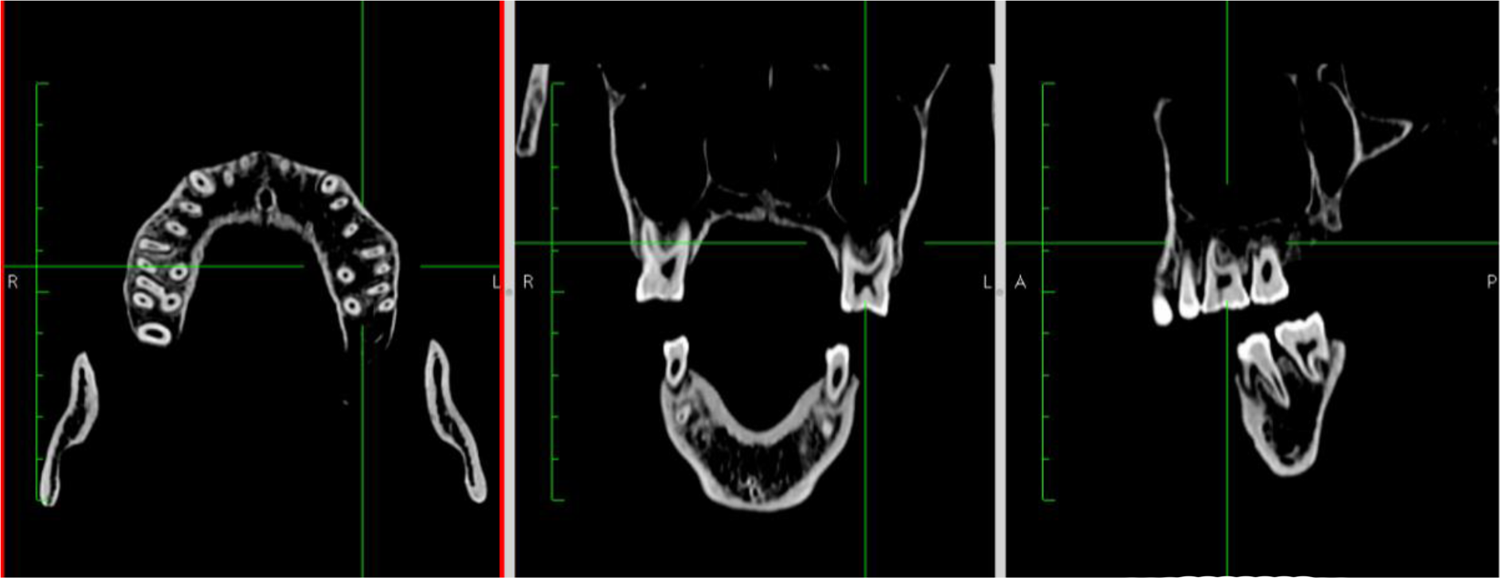
Horos Dicom Viewer 2D orthogonal view used to assess root and canal morphologies. **Left:** Coronal view at mid-point of roots. Centre: Anterior view at midpoint of roots. **Right:** Lateral view at midpoint of roots.

### Determination of root and canal number

Root and canal number are determined by applying the Turner Index (1991), which compares the point of bifurcation relative to total root or canal length, as measured in Horos Dicom Viewer from the cemento-enamel junction (CEJ) to the root apex/canal apical foramen (Figure 3). When this ratio is greater than one-third (33%) of the total root or canal length, the root or canal is classified as multi-rooted. When the ratio is less than one-third (33%) the root or canal is considered single rooted, or with a bifid apical third. Individual root/canal number for analysis is recorded as a simple numerical count (e.g., 1,2,3, etc.).

Using measurement tools in Horos Dicom Viewer, the midpoint of the root was measured halfway between the CEJ and the apices of the root/s (Figure 7). It is by these methods that data were acquired for the analyses described and carried out through this study. Here, a single root canal is defined as a canal which extends from the pulp chamber within the crown and exits at a single foramen. Accessory canals are not included in this study.

**Figure 7:**
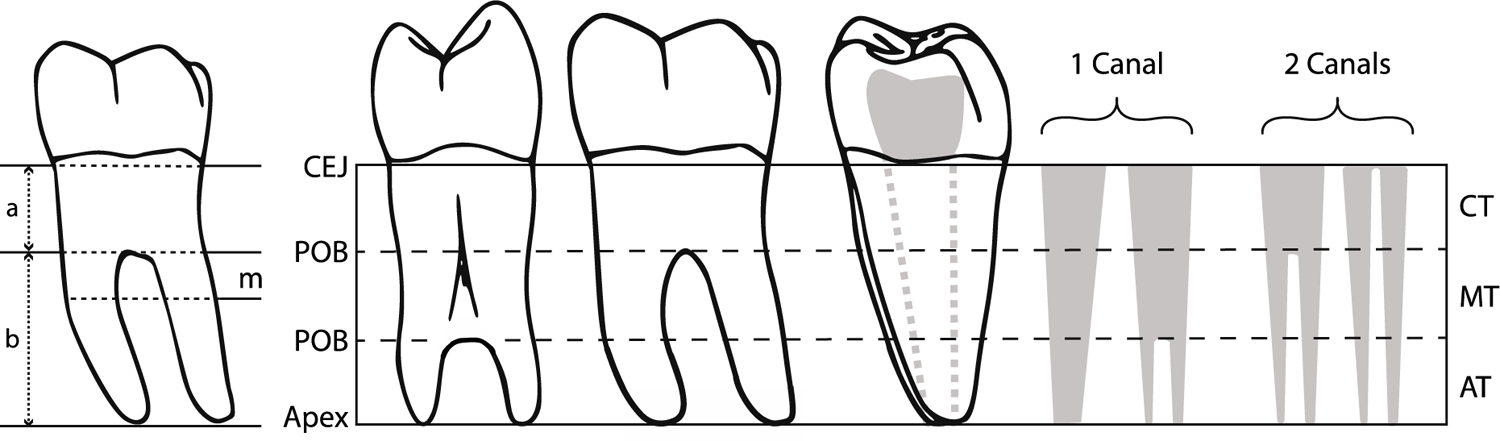
Left: Locations of measurements taken in Horos Dicom Viewer of (a) CEJ to point of bifurcation, (b) point of bifurcation to root apices, (m) mid-point between CEJ and root apices. **Right:** Determination of root and canal number. Distal view of single-rooted premolar with bifurcation of the apical third of the root. Lingual view of double-rooted mandibular molar. Distal root of double-rooted mandibular molar with examples of canal counts in solid grey. Dotted grey lines indicate canal/s position in root. CEJ = Cemento-enamel junction, POB = Point of bifurcation, Solid grey = canals. CT = cervical third, MT = middle third, AT = apical third.

### Statistical analyses

Data were analysed with the R Project for Statistical Computing. Because the osteological materials used in this study were recovered from excavation sites, many of the individuals comprising our sample are missing one or more teeth. As the mechanism causing these missing data are unrelated to the values of any variables used in analysis (missing completely at random), our observed values are essentially a random sample of the full data set and not biased (Sterne et al., 2009). Thus, multiple imputation is appropriate for our data set (Garson, 2015; Zhang et al., 2017). Using the missMDA package, we performed multiple imputation on missing data in preparation for analysis (Josse and Husson, 2016).

Because the Poisson distribution is typically used for count data, a Poisson general linear model (PGLM) was used to test the association between canal and root number at the *p* = 0.05 significance level (Zeileis et al., 2008). A key assumption underlying PGLM is the independence of observations (Hoffmann, 2004). Thus, the inclusion of multiple teeth from the same individuals may violate assumptions of independence for the PGLM used in this study. To account for this, we fit our PGLM with Generalized Estimating Equations (GEE).

GEE estimates population-averaged parameters and their standard errors based on a number of assumptions: (1) The responses variables are correlated or clustered; (2) There is a linear relationship between the covariates and a transformation of the response; and (3) within-subject covariance has a correlation structure (Zeger and Liang, 1986; Diggle et al., 2002). In order to determine our correlation structure and how root and canal number correlated within and between teeth we conducted a Pearson correlation analysis of canal and root number. We selected and Auto Regressive Order 1 (AR1) correlation structure for our GEE covariance matrix. While GEE estimates of model parameters are valid regardless of the specified correlation structure, the AR1 correlation structure is appropriate because it (a) has no distributional assumptions (Zuur et al., 2009); (b) can accurately model covariance for cross-sectional individual and clustered studies (Müller et al., 2009; Muoka et al., 2021); (c) accurately model within-subject correlation decreasing across time and/or space (Agresti, 2002); and (d) assumes observations within and individual are non-independent (Zeger and Liang, 1986). Thus, AR1 is appropriate at the individual and population levels, and for the temporospatial distances within and between individuals and groups within our sample. GEE was caried out using ‘geepack: Generalized Estimating Equation Package’ version 1.3.2 (Halekoh et al., 2006). Stepwise model selection was tested and quantified using Akaike Information Criteria (stepAIC). Tukey’s multiple comparison test was used for pair-wise analysis of population groups (Full statistical output is presented in Appendix Section 9.3). PGLM extended with GEE was also used to test for association between root and canal number by tooth and population groups. Tukey’s multiple comparison test was used for pair-wise analysis of population groups.

## Results

### Number of teeth, roots, and canals

Tables 3 and 4 report counts for number of roots and canals from post-canine teeth belonging to the right side of the maxilla and mandible. The number of roots in teeth from the sample are between one and four (Table 3). In this sample, teeth with four roots are limited to maxillary molars and appear with a relatively low frequency compared to 2 and 3 rooted teeth. Premolars, especially P_3_ and P_4_, are predominantly single-rooted, while the majority mandibular molars in this sample are double-rooted. Entomolaris (En), or three-rooted molars, appear in 18.05 % M1s, 1.23% of M2s, and 5.94% of M_3_s, and three-rooted paramolaris (Pa) appears in 3.63% of M_3_s.

**Table 3:**
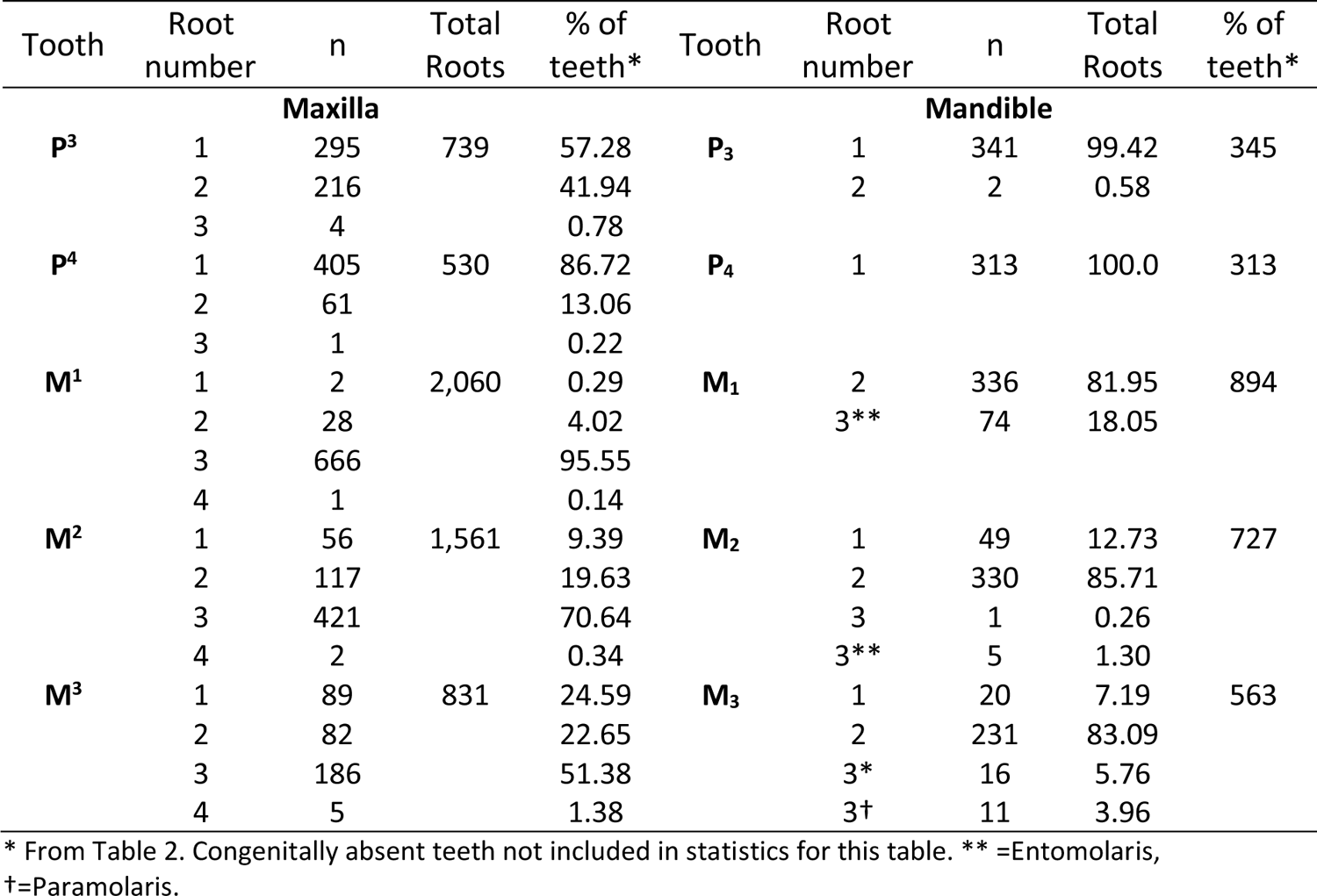
Number of roots in teeth of the maxilla and mandible by tooth

**Table 4:**
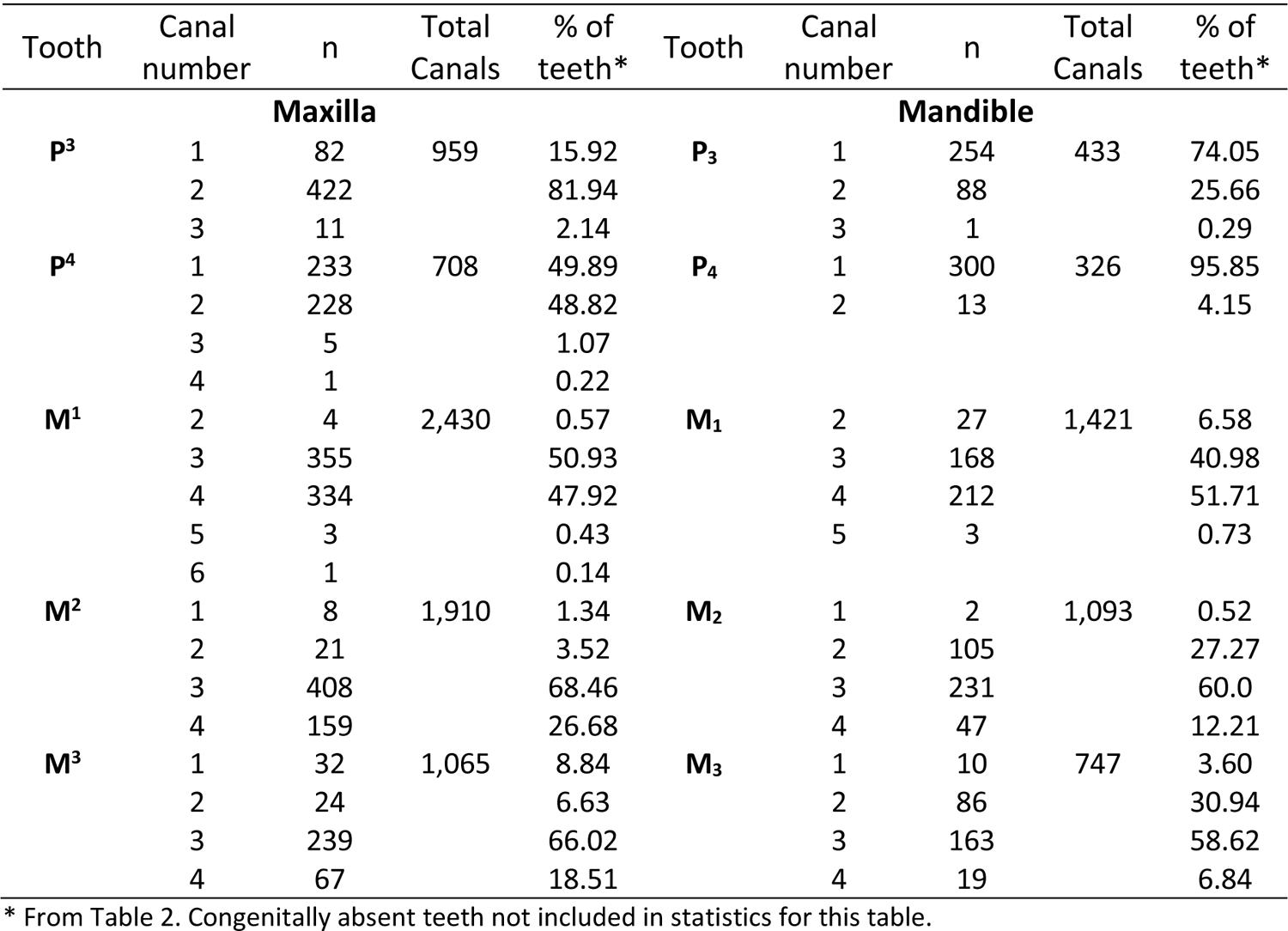
Number of canals per tooth in the maxilla and mandible by tooth

Teeth from this sample contain between one and six canals, and canal number often exceeds root number (Table 4). Many teeth contain two or more canals, especially in the molars. Molars have the greatest number of canals per tooth, with M^1^s showing the most variation in canal number.

### Inter-trait correlation

Pearson product-moment correlation coefficients (Figure 8) were computed to assess linear correlation between root number (RN) and canal number (CN) for teeth used in this study (Table 2). The majority of variables have negligible to weak positive or negative correlation coefficient strength values of 0.01 - ± 0.30 (Akoglu, 2018). Within same teeth moderate to strong correlation coefficient values of 0.31 - ± 0.69 (ibid) are found in P^4^ RN:P^4^ CN (0.46), M^3^ RN:M^3^ CN (0.47), M_2_ RN:M_2_ CN (0.35), and M_3_ RN:M_3_ CN (0.50). With the exception of P^3^ RN:P^4^ CN (0.46), P^3^ RN:P^4^ CN (0.65), P_3_ CN:P_4_ CN (0.43), M^3^ RN:M^2^ CN (0.31), and M_2_ CN:M_3_ CN (0.31), there are no significant correlations of RN to CN across different teeth.

**Figure 8:**
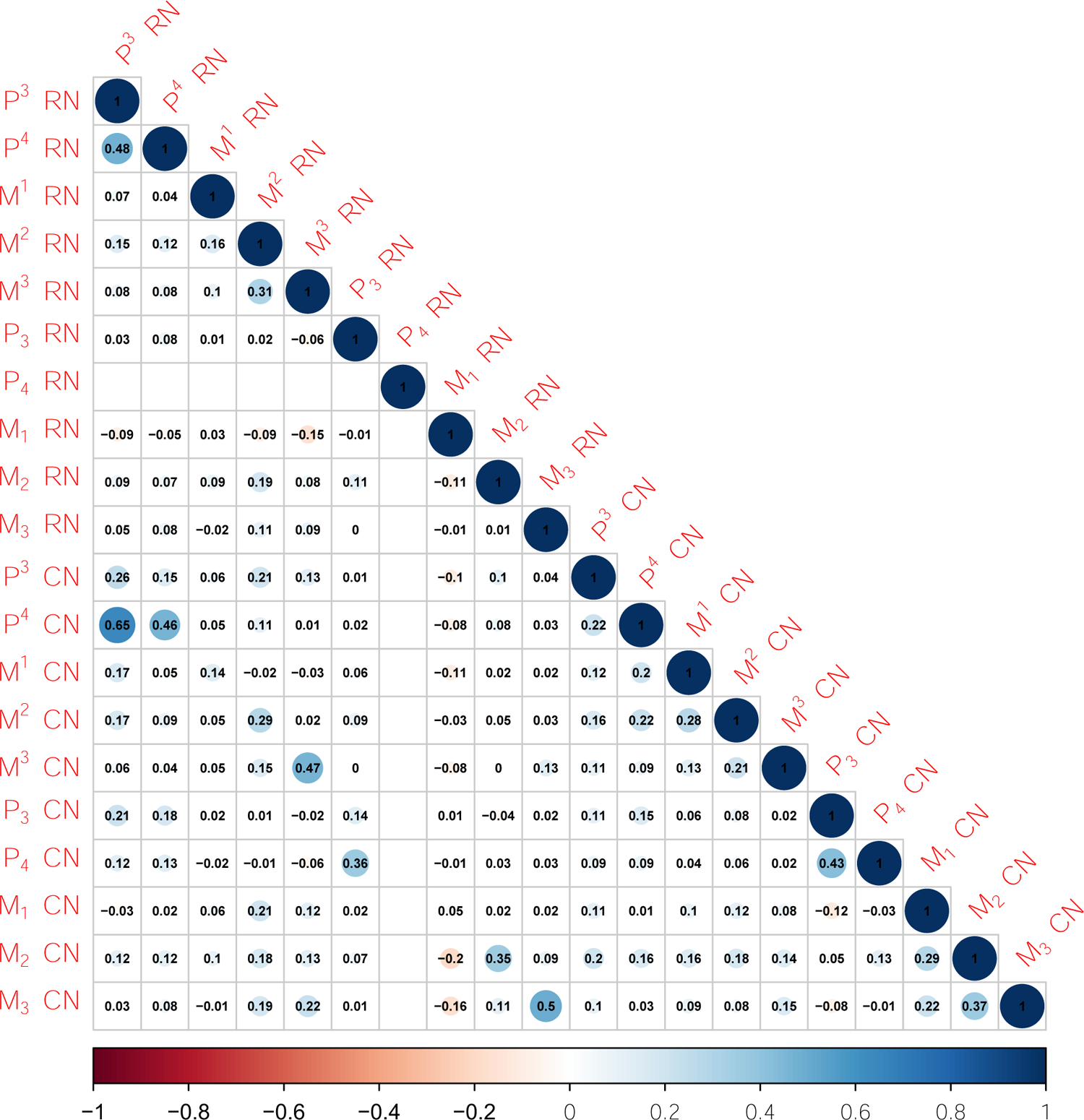
Pearson correlation of root number (RN) to canal number (CN). Significance level = 0.05. Significant positive correlation coefficients in blue. Significant negative correlation coefficients in red. Blank cells in P4 RN:P4 RN due to all P4s having the same level (i.e., one root. See Table 3).

### PGLM of relationship between canal and root number in individual teeth

While independent variables are uncorrelated, uncorrelated variables are not always independent. To address this, we fit PGLM with GEE to account for low levels of correlation between some traits (Figure 8), and to account for using multiple teeth from the same individuals, which may violate assumptions of variable independence. PGLM fitted with PGEE was used to directly test the linear relationship of root to canal number by tooth - in other words, to see how the relationship between canal and root number varies across different tooth types. PGLM of individual teeth reveal that for M^1^-M^3^, and M_1_-M_3_, as canal count increases, so does root count (Table 5). In the maxilla, the greatest increase in root to canal number is found in M^1^ (99.99%), and similar relationships are found in M^2^ and M^3^.

**Table 5.** Regression parameters for PGLM and GEE regression of the association between canal-to-root number by tooth, ranked by odds ratios from greatest to least*

|  | Estimate | Odds ratio | Std. error | Wald | P-value |
| --- | --- | --- | --- | --- | --- |
| *Canal number* | 0.117 | 1.124 | 0.007 | 251.300 | < .0001 |
| **Maxilla** |  |  |  |  |  |
| M<sup>1</sup> | 0.693 | 1.999 | 0.025 | 750.400 | < .0001 |
| M<sup>3</sup> | 0.650 | 1.916 | 0.025 | 683.700 | < .0001 |
| M<sup>2</sup> | 0.648 | 1.911 | 0.025 | 651.900 | < .0001 |
| P<sup>3</sup> | 0.000 | 1.000 | 0.018 | 0.000 | 0.990 |
| P<sup>4</sup> | -0.091 | 0.913 | 0.011 | 69.100 | < .0001 |
| **Mandible** |  |  |  |  |  |
| M<sub>3</sub> | 0.356 | 1.428 | 0.022 | 264.800 | < .0001 |
| M<sub>2</sub> | 0.127 | 1.135 | 0.049 | 2.604 | < .0001 |
| M<sub>1</sub> | 0.287 | 1.330 | 0.028 | 106.700 | < .0001 |
| P<sub>4</sub> | -0.119 | 0.887 | 0.008 | 248.000 | < .0001 |
| P<sub>3</sub> | -0.127 | 0.881 | 0.008 | 236.100 | < .0001 |

\*Model fitted without intercept.

Maxillary premolars remain relatively stable, with a minimal increase (0.03%) in P^3^, and no increase in root number in P^4^. Mandibular molar (M_1_-M_3_) roots are comparatively similar to one another in their odds ratios, especially M_1_ and M_2_; while surprisingly, mandibular premolars (P_3_-P_4_) show that as canal number increases root number does not

Prediction curves differ for each tooth, and the maxilla and mandible as a whole (Figure 9). Similar tooth groups have similar prediction curves — P_3_, P_4_, and P^4^; M_3_, M_2_, and M_1_; and M^1^, M^2^, M^3^; and these differ between the maxilla and mandible. As prediction curves diverge from the 1:1 canal-to-root ratio, root number decreases, though this does not necessitate a concomitant reduction in canal number (Figure 5).

**Figure 9:**
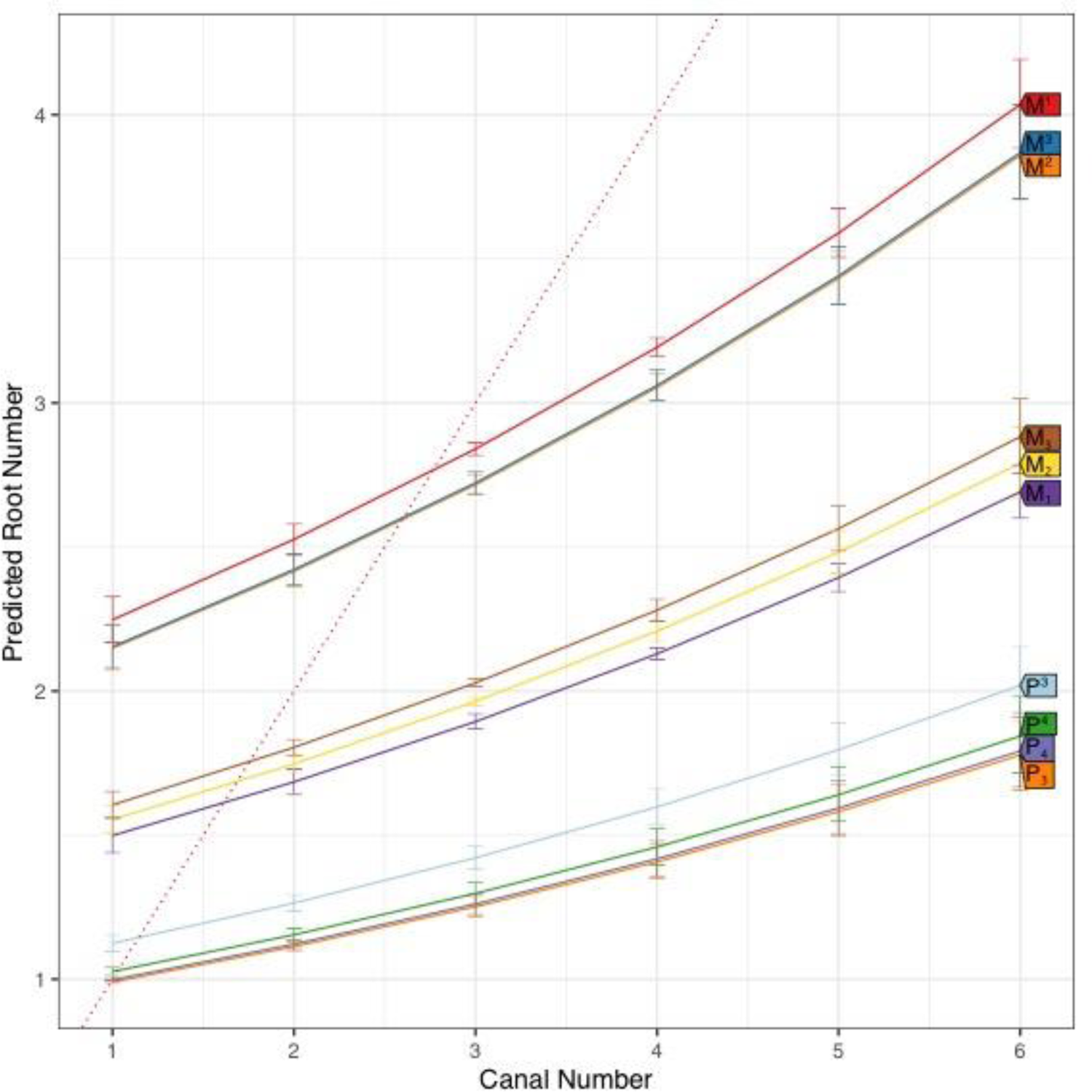
PGLM prediction curves with error bars for canal to root number for individual teeth. Dotted red line represents 1:1 canal to root relationship (i.e., what would be observed if there was a simple 1:1 relationship between roots and canals). Over prediction in the number of roots for single canaled M^1^/M1s-M^3^/M3s is owing to very small sample of individuals with one root to one canal (see Table 3 for counts).

Figure 10 plots proportions of canal to root number of individual teeth within the sample. Different patterns are clearly evident across all teeth and between the maxilla and mandible and help to explain groupings of individual tooth prediction curves in Figure 9.

**Figure 10:**
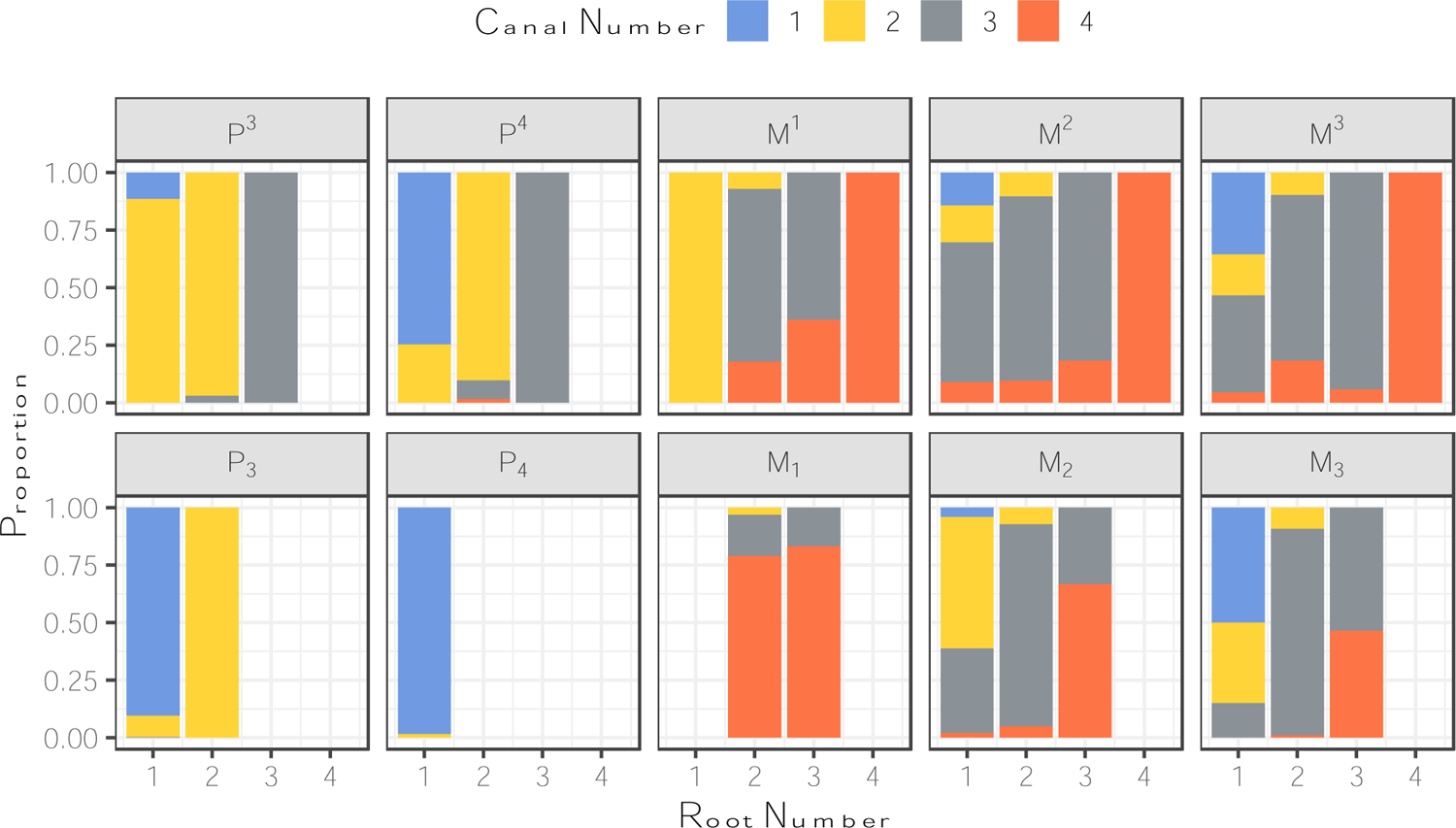
Proportion of canal to root number for individual teeth. 5 canaled teeth = 14, 6 canaled teeth = 2, are included in calculations of proportions but are not visualized on this plot due to small sample size.

There is a slight over-prediction in the number of roots for single canaled M^1^-M^3^’s owing to 1) very small sample of individuals with one root to one canal for these teeth (see Table 3 for counts); and 2) because we have used a fixed non-parametric model to capture the non-linearity between canal and root number. Variation in canal to root number decreases in the premolars while increasing in the molars, though this variation does not covary between opposing individual maxillary and mandibular teeth. The greatest variation is found in the maxillary molars (M^1^-M^3^) while the least in found in P_4_.

Tukey pair-wise comparisons of PGLM of root to canal number by tooth (Figure 11) show that patterns in prediction curves and canal-to-root proportions plotted in Figures 9 and 10 reflect significant differences between teeth. Full statistical output is presented in the Supplementary Materials Table B.

**Figure 11:**
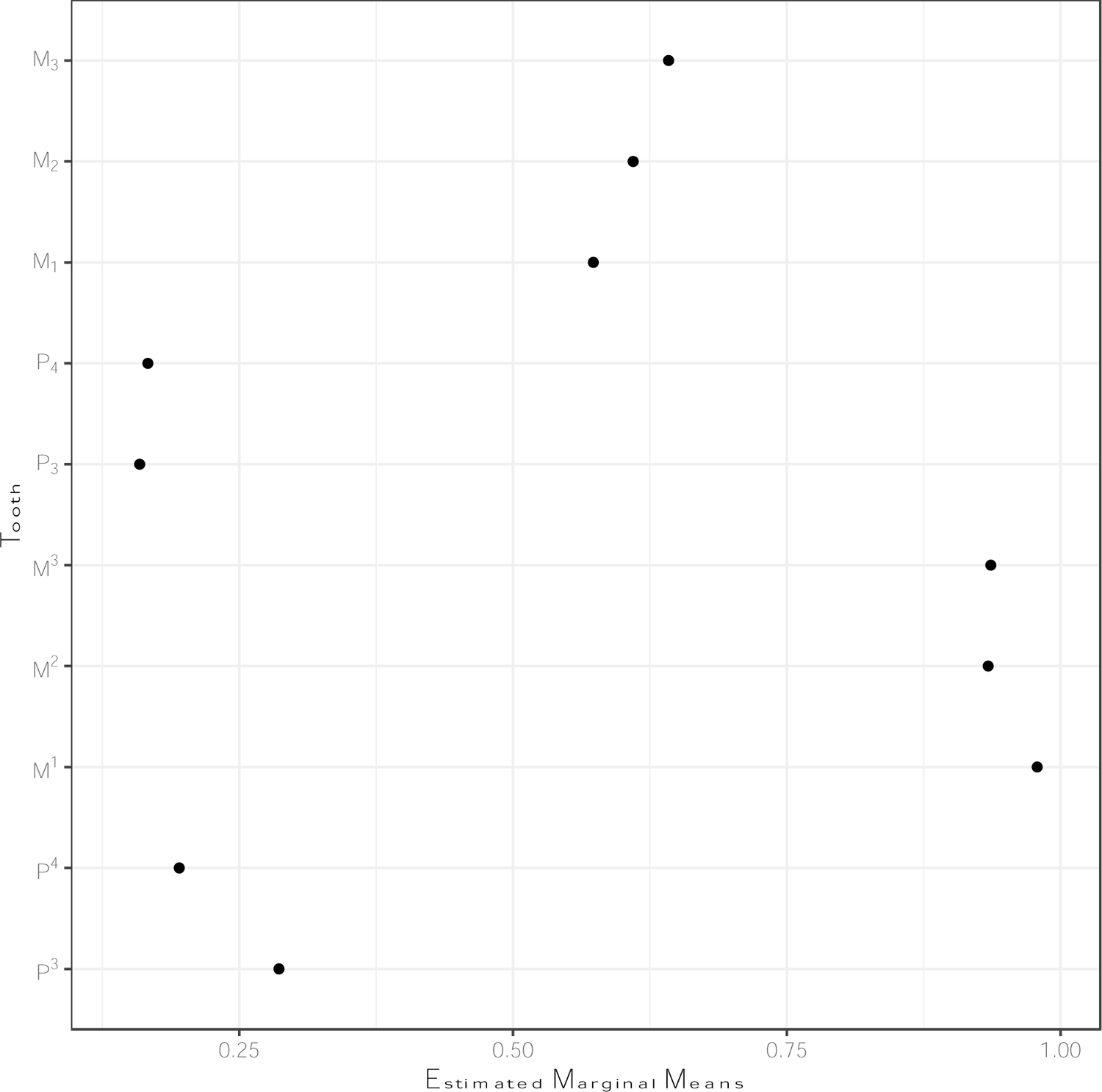
Estimated marginal means derived from Tukey pair-wise comparisons of canal to root number by tooth. Black dot= mean value; Blue bar = confidence intervals. The degree to which red comparison arrows overlap reflects the significance (*p* = 0.05) of the comparison of the two estimates. Full statistical output is presented in Supplementary Materials Table B.

### PGLM of relationship between canal and root number in geographical populations

We used PGLM to test the linear relationship of root to canal number by tooth across population groups (Table 6). To avoid emphasizing results against one geographical region or tooth we fitted the model without an intercept.

**Table 6:** Regression parameters for the PGLM testing the association between canal and root number by tooth in geographic populations, ranked by odds ratio from greatest to least*.

| Groups | Estimate | Odds ratio | Std. error | Wald | P-value |
| --- | --- | --- | --- | --- | --- |
| Canal number | 0.329 | 1.389 | 0.002 | 29139.9 | < .0001 |
| Sub-Saharan Africa | -0.211 | 0.810 | 0.007 | 816.8 | < .0001 |
| Sino-Americas | -0.235 | 0.790 | 0.006 | 1749.3 | < .0001 |
| Sunda-Pacific | -0.250 | 0.779 | 0.010 | 658.8 | < .0001 |
| West Eurasia | -0.238 | 0.789 | 0.007 | 1065.3 | < .0001 |
| Sahul-Pacific | -0.258 | 0.773 | 0.007 | 1206.2 | < .0001 |

\* Model fitted without intercept

Individual teeth of geographical groups are relatively similar in their odds ratios and prediction curves; and follow a similar pattern of divergence from a 1:1 canal-to-root ratio (Figure 12). Prediction curves for Sub-Saharan African populations are closest to the 1:1 canal-to-root ratio, while Sino-American populations are the furthest. While odds ratios of Sub-Saharan African and Sino-American populations are most similar, marginal effects quantify how both groups vary differently in their canal to root ratios (Figure 13).

**Figure 12:**
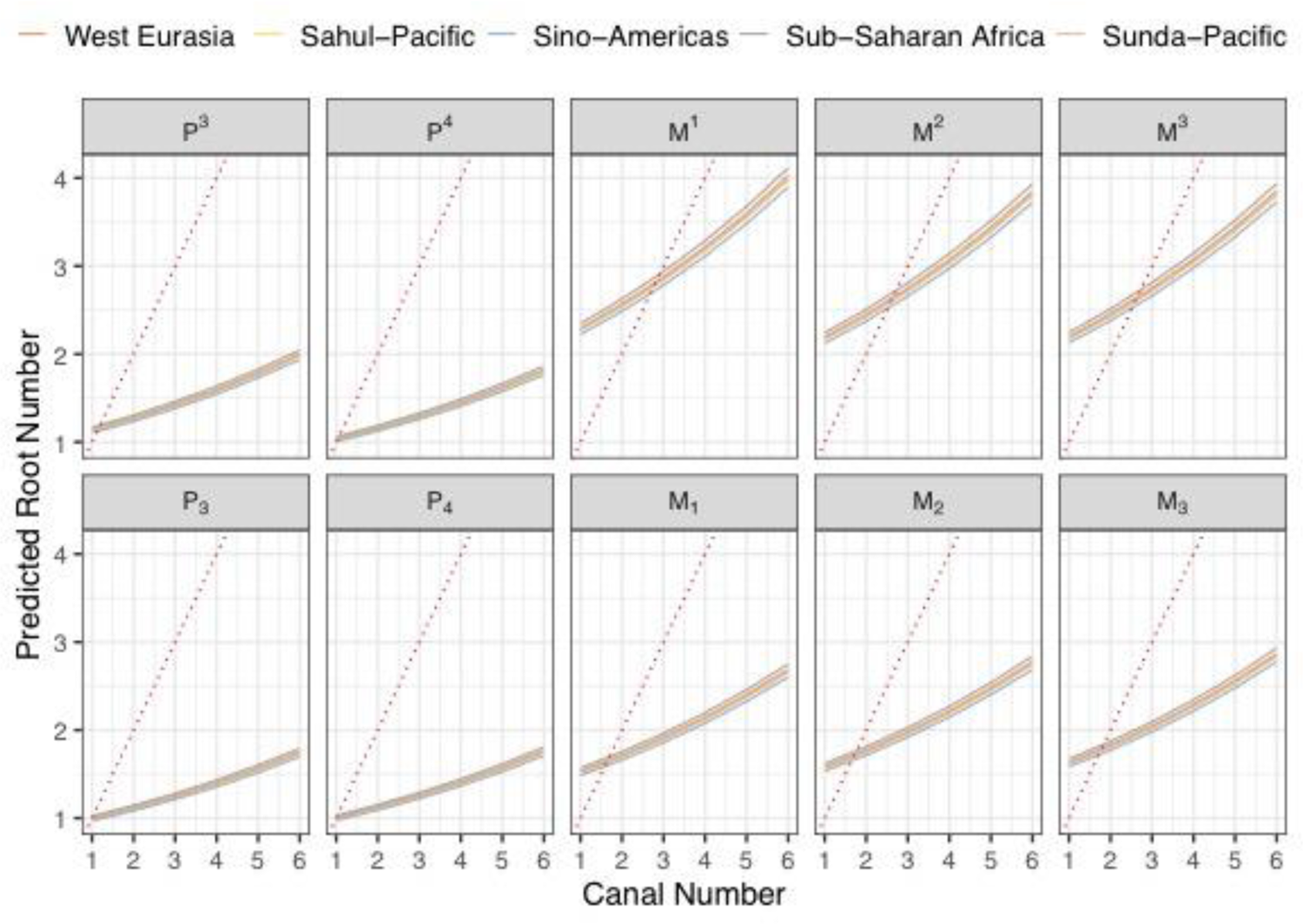
PGLM prediction curve for root to canal number by population. Dotted red line represents 1:1 root to canal relationship. Over prediction in the number of roots for single canaled M^1^/M1s-M^3^/M3s is owing to very small sample of individuals with one root to one canal (see Table 3 for counts).

**Figure 13:**
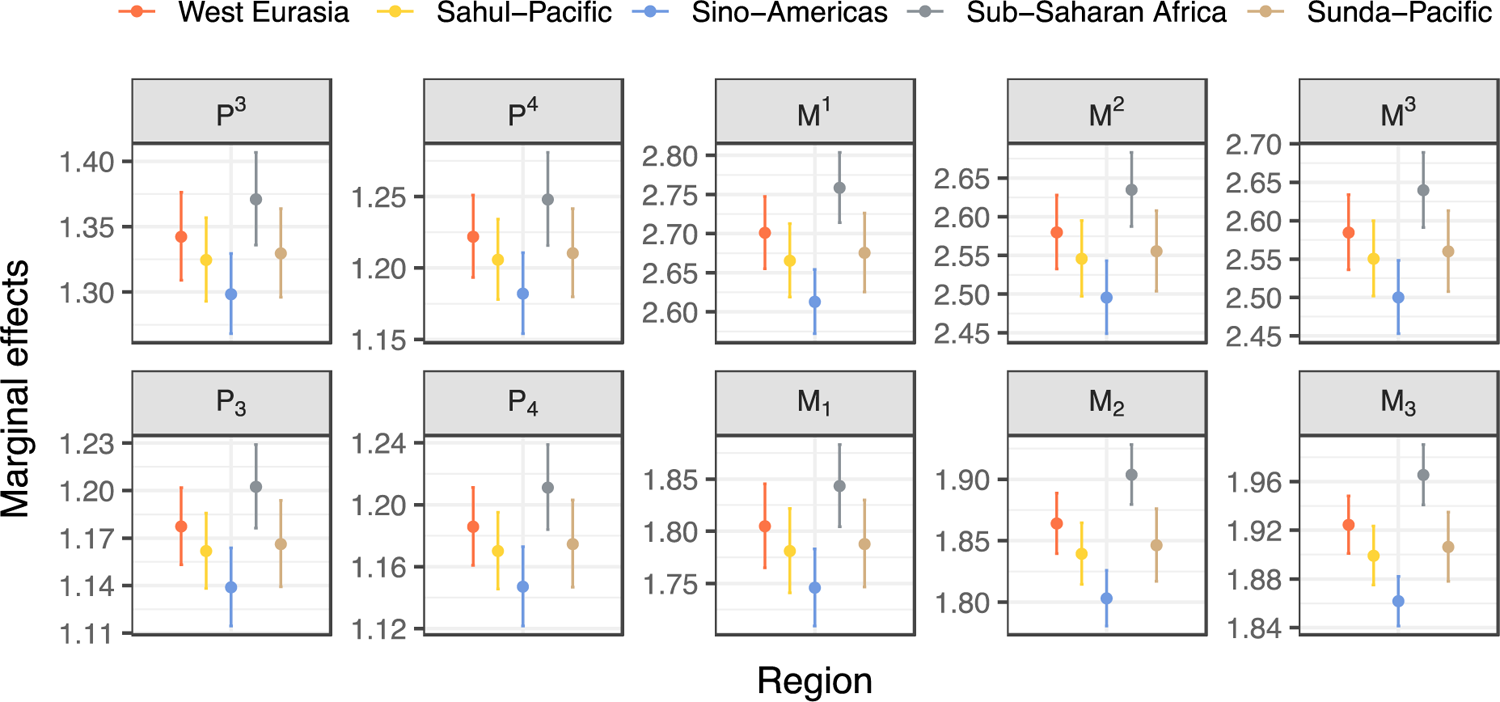
Marginal effects of canal to root count in individual teeth by geographical region.

Marginal effects quantify how both groups vary differently in their canal to root ratios (Figure 13) when the explanatory variable (canals) changes by one unit. For all teeth, the Sino-American groups have the lowest percentage of change in root number as canal number increases, while Sub-Saharan Africans show a higher percentage of root number change as canal number increases.

Tukey pair-wise comparisons of PGLM of canal to root number by population show that patterns in prediction curves and marginal effects plotted in Figures 12 and 13 reflect significant differences between Sub-Saharan Africa and all other groups (Figure 14). Significant differences are also shown between Sahul-Pacific and Sino-Americans. Full statistical out is provided in Supplementary Materials Table C).

**Figure 14:**
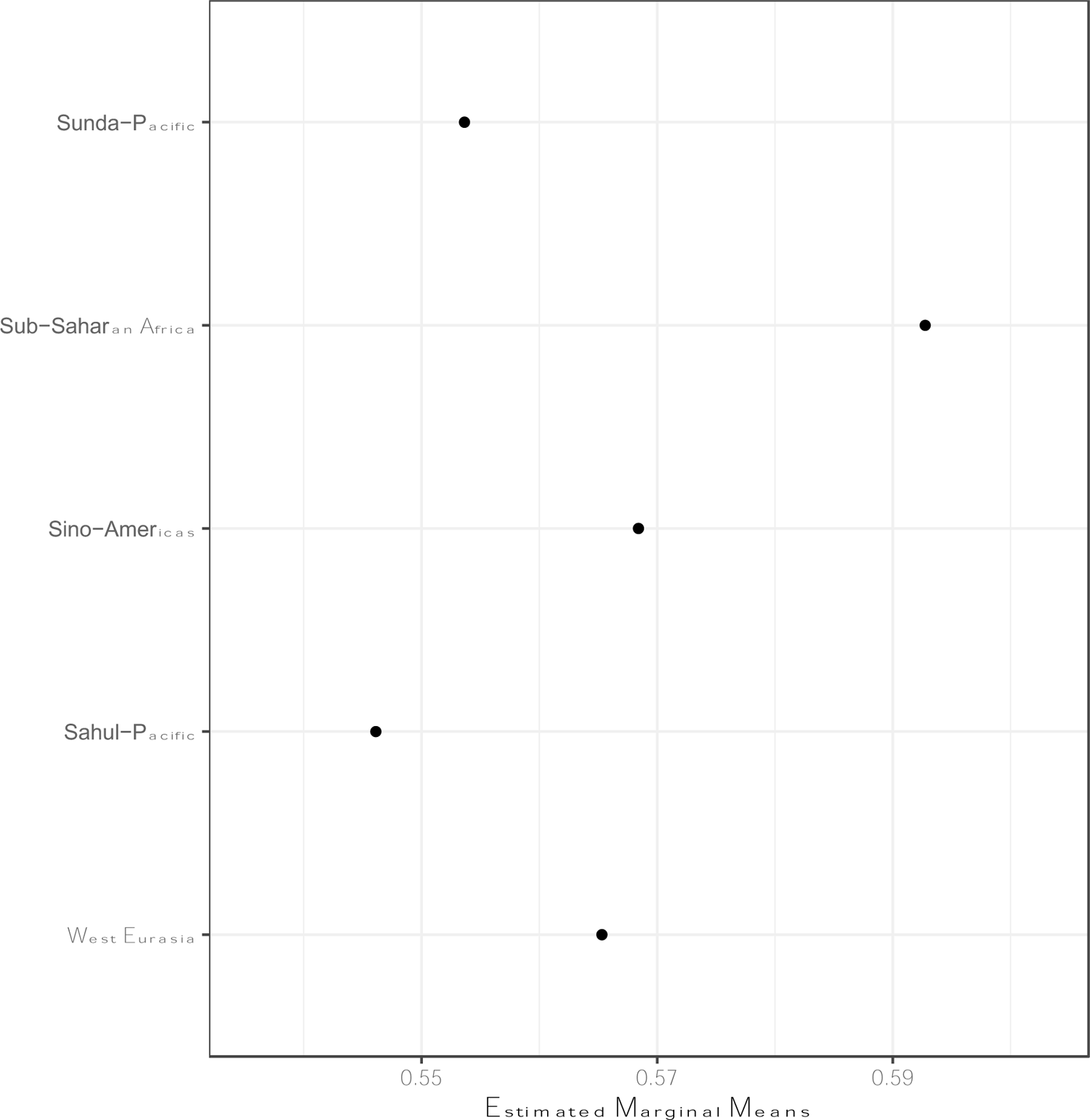
Estimated marginal means derived from Tukey pair-wise comparisons of PGLM of canal to root number by geographical region. Black dot= mean value; Blue bar = confidence intervals. The degree to which red comparison arrows overlap reflects the significance (*p* = 0.05) of the comparison of the two estimates. Full statistical output is presented in Supplementary Materials Table C.

## Discussion

In the analyses presented above we have been able to show that there is little to no correlation between root and canal number in teeth (Figure 8). However, because uncorrelated random variables are not always independent, we extended our PGLM with GEE to develop a predictive model of the relationship between canal and root number, globally and by region, and we show that this relationship is not perfectly linear. We have found that canal number predicts root number, and that the greater the number of canals the more complex, and less predictable the number of roots. This relationship varies by maxillary and mandibular teeth and tooth row. These results raise a number of issues: what does the complexity of canal to root number relationships mean developmentally? Why does this complexity vary across particular tooth types? How does divergence in canal and root number vary between maxillary and mandibular teeth in total, by population, and individually?

### Differences in root and canal number

Currently, there is no consensus as to why canals and roots should differ in number, given that canal formation precedes root formation. Clusters of blood vessels entering the dental papilla early in tooth formation coincide with the positions where roots will eventually form (Nanci, 2012). The HERS and expanding dental pulp form around these nerves and blood vessels before dentin formation. Thus, each root must contain at least one canal for the pulp, and the nerve and blood supply that precede the formation of the surrounding root structure. It is possible that number, size, and configuration of blood and nerve supplies is, in part, responsible for variation in canal number with the roots, and not variation in the number and orientation of the interradicular processes alone.

### Variance across teeth and between the maxilla and mandible

Why canal and root number should vary both within and between teeth of the maxilla and mandible is also unknown. Prediction curves and proportions of canal to root numbers show that the relationship between canals and roots within tooth types are similar to one another, i.e., maxillary molars are alike, while being significantly different from other tooth types, i.e., such as premolars and mandibular molars. Similar estimates (Table 5) and PGLM curves of tooth types (Figure 9) seem to lend support to the morphogenetic field model in which teeth within a field are more similar to one another than to teeth of another field (Butler, 1937, 1963; Dahlberg, 1945); especially for molar fields in both jaws. These results suggest that the number of canals and roots within tooth types are relatively “fixed” with little intra-tooth type variation. We propose two possible explanations, the first functional, the second spatial.

Megadonty is a hallmark of early hominin evolution (Robinson, 1956; Wood and Abbott, 1983; Reed, 1997; Wood and Constantino, 2007); and heavy chewing requires large teeth. The majority of chewing actions occur on the broad occlusal surfaces of the post-canine teeth where, compared to anterior teeth, masticatory movements are complex combinations of antero-posterior, vertical and lateral movements (van Eijden, 1991; Ledogar et al., 2016). Chewing pressures on the maxillary teeth result from absorption of shearing and compressive forces generated by the active movement of the mandible (Ledogar et al., 2016). During mastication, maxillary molars are subjected to greater medio-lateral directed loads than mandibular molars (Dempster et al., 1963; Spears and Macho, 1998). These medio-lateral forces are dissipated into the jaws via the tooth roots (Nikolai and Zwemer, 1985; Baragar and Osborn, 1987); and, in humans, are strongest at, and decrease posteriorly from M^1^/M_1_s (Gordon, 1984; Macho and Spears, 1999). Consequently, as root surface area decreases in M^2^ and M^3^, so does root number ((Dempster et al., 1963); Table 3).

It is possible that where increased masticatory loadings are a selective pressure for larger teeth, an increased blood supply required for developing a larger tooth will result in an increase in canal number. This will, in turn, result in more roots. The increased mesio-distal and bucco-lingual dimensions of premolars tooth crowns belonging to megadontic “robust australopiths” (*Paranthropus boisei*, *P. robustus*, *P. aethiopicus*), support such as hypothesis. These “hyper-robust” hominins regularly had multi-rooted/canaled premolars (Robinson, 1954, 1956; Wood et al., 1988b; Brook et al., 2014; Moore et al., 2016; Kupczik et al., 2018), and the ancestral hominin phenotype has been proposed as three-root maxillary premolars, and two-root mandibular premolars In modern humans, molars withstand the heaviest masticatory loadings while premolars are subjected to the least (Demes and Creel, 1988; Ledogar et al., 2016). That masticatory stresses produce high strains in the alveolar margin of the anterior maxilla (Ledogar et al., 2016) may act to increase canal and root number in the maxillary premolars compared to mandibular premolars. Developmentally, Shields (2005) proposed that tooth germ size influenced the number and development of IRP’s. However, multiple studies have noted that tooth crown size (used as proxy for tooth germ size) does not always covary with root number and size in humans and hominoids (Abbot, 1984; Shields, 2005; Moore et al., 2013, 2016).

Different masticatory forces resulting from dietary demands have been shown to increase tooth root surface area, and thus size, in primates (Kovacs, 1971; Spencer, 2003; Kupczik and Dean, 2008; Ledogar et al., 2016). A possible selective mechanism to increase tooth root surface area would be to increase the number of roots, which would in turn enlarge the cervical base area of the crown (Kupczik et al., 2005). A study of *Gorilla gorilla*, *Pan troglodytes*, as well as of 26 fossil gracile and robust hominins from South Africa concluded that dietary adaptations produced mesio-distal expansion at the base of tooth roots in M^1^s (Kupczik et al., 2018). The authors (ibid) concluded that it was increases in root splay that accommodated higher masticatory loadings, but that the mesio-distal expansion of the root bases in robust hominins might be an adaptive response to different jaw kinematics for chewing different food types — horizontally directed repetitive chewing in *P. boisei* (Demes and Creel, 1988; Wood and Constantino, 2007), versus multi-directional loading of *P. robustus* (Macho, 2015). However, the extant and fossil species from this study are already characterized by multi rooted molars and premolars (Sperber, 1973; Wood et al., 1988a; Kupczik et al., 2005; Shields, 2005); so it is difficult to discern if mesio-distal expansion of the roots is an adaptive response to biomechanical pressures, a bi-product of additional roots, or both. If root splay is in fact the primary adaptive response to increased masticatory loading, the selective pressures underlying what point single root surface area/size stops increasing and root differentiation begins have yet to be elucidated.

Alternatively, variation may arise from space required for growing teeth in the developing jaws. Consider that maxillary and mandibular 1st molars are the first adult teeth to erupt (at 6-7 years) followed by the anterior teeth (7-10 years), premolars (10-12 years), followed by 2nd (12-13 years) and 3rd molars (17-21 years). In this spatial scenario maxillary and mandibular 1st molars have the greatest number of roots and canals, while late-forming and erupting premolars have the least as they are sandwiched between 1st molars and the already erupted anterior teeth. Constrained variation, especially in the premolars may be explained by limited space for growth and development, while maxillary and mandibular molars have spatial restrictions on their growth and development limited by dimensions of the palate and by the ascending ramus of the mandible.

Biomechanical and spatial explanations need not be mutually exclusive. It may be the case that canal, and root variation found in modern humans is a product of reduction in space as a consequence of reduced selection for intensive biomechanical chewing pressures in early human evolutionary history. Premolar root number has been documented as more variable than in all other tooth types (Sperber, 1973; Wood et al., 1988a; Kupczik et al., 2005; Shields, 2005). Contrary to the molarization of the robust Paranthropines, the reduction of premolar root number is present in South-African gracile hominins. Robinson (1956) and Sperber (1974) report predominantly (84%) double-rooted maxillary premolars in a sample of *Australopithecus africanus*, though single (8%) and triple-rooted (8%) variants do occur. *A. africanus* mandibular premolars are reported as having single C-shaped (also referred to as Tomes’ root) and double-rooted mandibular molars (Robinson 1952, 1956; Sperber, 1974, Moore et al.,2016). Thus, this trend for reduction in premolar root number appears early in human evolutionary history (3.4 - 2.4 Ma) and coincides with dietary shifts towards meat and/or softer cooked foods (Luca et al., 2010), and reduction of hominin tooth crowns, jaws and face. At 1.8 Ma *Homo erectus* has fewer tooth roots, especially M_3_’s, than earlier members of our genus, and *H. erectus* premolars are frequently single rooted (Antón, 2003). This trend in root number reduction continues through more recent members of Genus *Homo* including some specimens allocated to *H. heidelbergensis* and *H. neanderthalensis* (Benazzi et al., 2011; Martinón-Torres et al., 2012; Zanolli and Mazurier, 2013).

### Differences in geographical groups

Sino-American and Sub-Saharan African populations are significantly different from one another (Figure 11, Supplementary Materials Table C), and these differences can be explained by reduction in root number in the former. Compared to all other groups, Sino-Americans have the greatest proportion of single rooted teeth across all populations while Sub-Saharan African populations have the smallest (Figure 10). This trend is present regardless of canal number. The exception to this is the presence of three-rooted mandibular molars (ento- and para-molaris root forms) in Sino-Americans (Carlsen and Alexandersen, 1990). This form represents a relatively rare polymorphism, and appears with frequencies around 30-50% in East Asian, Inuit, and Aleut populations; 5-15% in Southeast Asian and Pacific populations; compared to 1% in European and Sub-Saharan African populations (Scott et al., 2018).

As with tooth types, there is no clear explanation for changes in canal to root number between populations. The reasons may be biomechanical in nature and relate to different diets between populations. However, this is unlikely as the Sino-American populations included in this study (primarily comprised of North American First peoples; see Figure 4 & Supplementary Materials Table C), did not pursue uniform subsistence strategies. However, the effect of different diets on tooth root and canal morphologies is poorly understood, with only a few studies centred on non-human primates, and gracile and robust Australopiths (see Kupczik et al., 2018 for an overview).

The study of dental traits have an extensive history and utility for characterizing and assessing the biological relationships within and populations (see Scott et al., 2018 for a comprhensive review). Dental morphology has been shown to be under strong genetic control and minimally affected by environmental factors (Corruccini et al., 1986; Dempsey and Townsend, 2001). The evolutionary trend of teeth has also been described as towards reduction in size and simplification in morphology (Scott and Turner II, 1988). While the authors of these studies were describing tooth crowns, tooth roots are presumably operating under the same genetic and environmental constraints, and evolutionary trends.

PGLM predictions (Table 6) and marginal effects (Figure 13) support evidence of simplification in terms of reduction. Sub-Saharan Africans and Sino-Americans are furthest in distance from one another in time and space, and the former group shows the greatest variation in root and canal number, while the latter shows a reduction. For example, Sino-Americans have a higher proportion of single rooted, double-canaled M_2_’s and M_3_’s than all other groups. Additionally, congenitally absent M_3_’s are common (>25%) in Sino-American populations (Turner II et al., 1991; Daito et al., 1992; Rakhshan, 2015; Scott et al., 2018).

Compared to Sub-Saharan African populations, Western Eurasia, Sahul- and Sunda-Pacific groups have reduced variability, though not as much as Sino-Americans. These three groups share similar linear relationships (Figure 8) and canal to root proportions (Figure 10), though marginal means of West Eurasian and Sunda-Pacific populations reveal their canal to root relationships are more similar to Sub-Sharan Africa, while Sahul-Pacific is closer to Sino-Americas.

Recent studies have highlighted the decrease of genetic and phenotypic diversity in human populations with increasing distance from Sub-Saharan Africa (Handley et al., 2007). This decrease in diversity has been interpreted as evidence of an African origin for anatomically modern humans. Reduced intra-population diversity has been ascribed to an “Out of Africa” migration, and sequence of founder events due to rapid expansions and colonization of the world (Prugnolle et al., 2005; Liu et al., 2006; Li et al., 2008). This reduction in diversity has been recorded in human dental (Hanihara and Ishida, 2005; Hanihara, 2008), craniofacial (Hanihara and Ishida, 2009; Betti et al., 2012), and morphometric traits (Manica et al., 2007), further supporting genetic hypotheses of this single, African origin and subsequent expansions. However, some exceptions to this exist. For example, three rooted M_1_’s, sometime referred to as Radix entomolaris (see Calberson, De Moor and Deroose, 2007 for a review), increase in Sino-American populations while appearing in low frequency in other populations; especially Sub-Saharan Africa (Scott et al., 2018). This trait has been most commonly attributed to genetic drift (Scott et al., 2018), though a recent study has suggested archaic introgression (Bailey et al., 2019); however, see Richard Scott, Irish and Martinón-Torres (2020) for a rebuttal.

## Conclusion

This paper presents a novel investigation into the relationship between canal and root number in human post-canine teeth. In all cases, canal number is either equal to or exceeds root number, supporting our hypothesis that canal number precedes and is, in part, responsible for root number in all post-canine teeth. These canal to root relationships are significantly different between tooth types (i.e., molars and premolars), within and between the maxilla and mandible. Results indicate that Sub-Sharan African and Sino-American populations are significantly different in their canal to root numbers, and this difference represents an overall reduction in root number with distance from Africa, but not necessarily canal number. Canal to root relationships differ across all populations studied, however the reasons for these differences are not ultimately clear. To test population affinities and differences, future studies should include morphological distance-based analysis to test divergence, as well as consider additional biological, historical, linguistic and cultural data. Results also show that tooth types within and between the jaws have different linear relationships and that these relationships are significantly different. These results support biomechanical and spatial hypotheses related to tooth crown size in hominin evolution, and future studies should include root *and* canal count in their analysis.

## Acknowledgements

We would like to thank Professor Marta Miraźon-Lahr and Dr. Frances Rivera for permitting use of their CT scans from the Duckworth Collection at the Leverhulme Centre for Human Evolutionary Studies, University of Cambridge; and Dr. Lynn Copes for permitting use of her CT scans, collected for her PhD dissertation, from the Smithsonian National Museum of Natural History and American Museum of Natural History, and the editors and reviewers for their feedback. We thank the Duckworth Laboratory, University of Cambridge, for permission to study material within its collections. We would also like to thank Dr. Christopher N. Foley for assistance and feedback on our statistical analyses; and Friederike Jürcke for assistance with Figures 2 & 5.

## Author contributions

Concept and design by Jason Gellis and Robert Foley. Acquisition of data and data analysis by Jason Gellis. Drafting of the manuscript by Jason Gellis and Robert Foley. Revision of the manuscript by Jason Gellis.

## Supplementary Material

**Table A:**
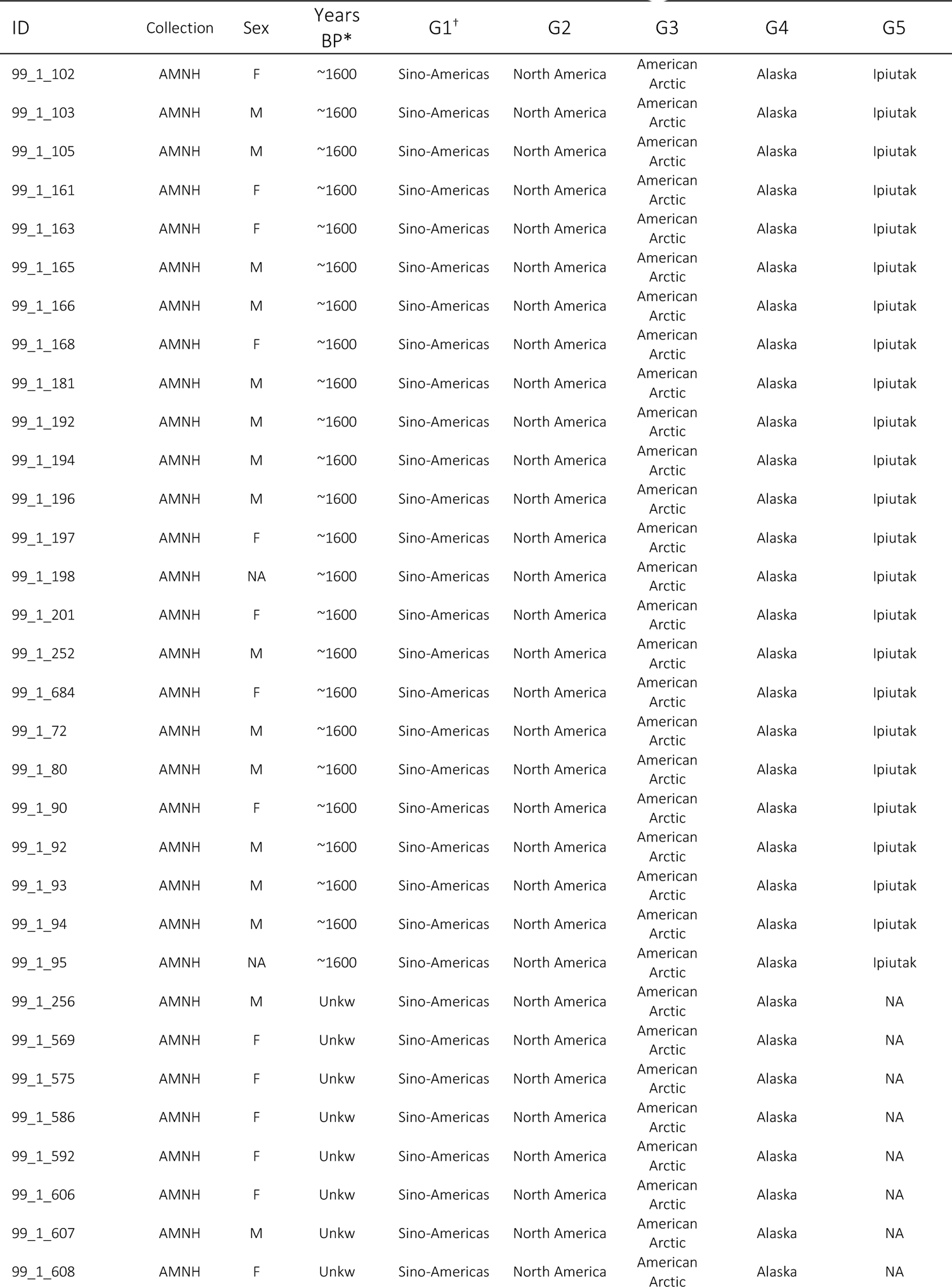
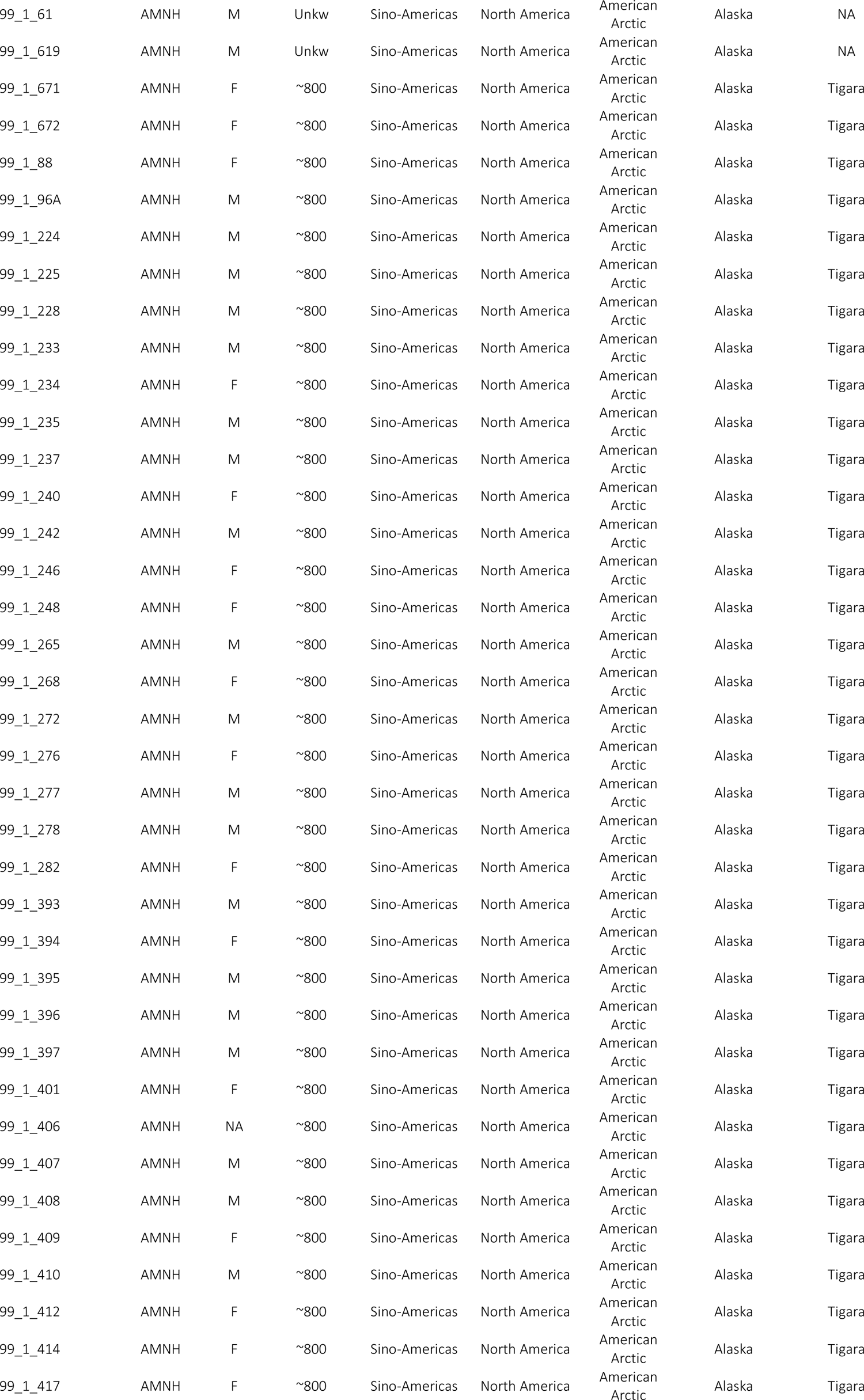
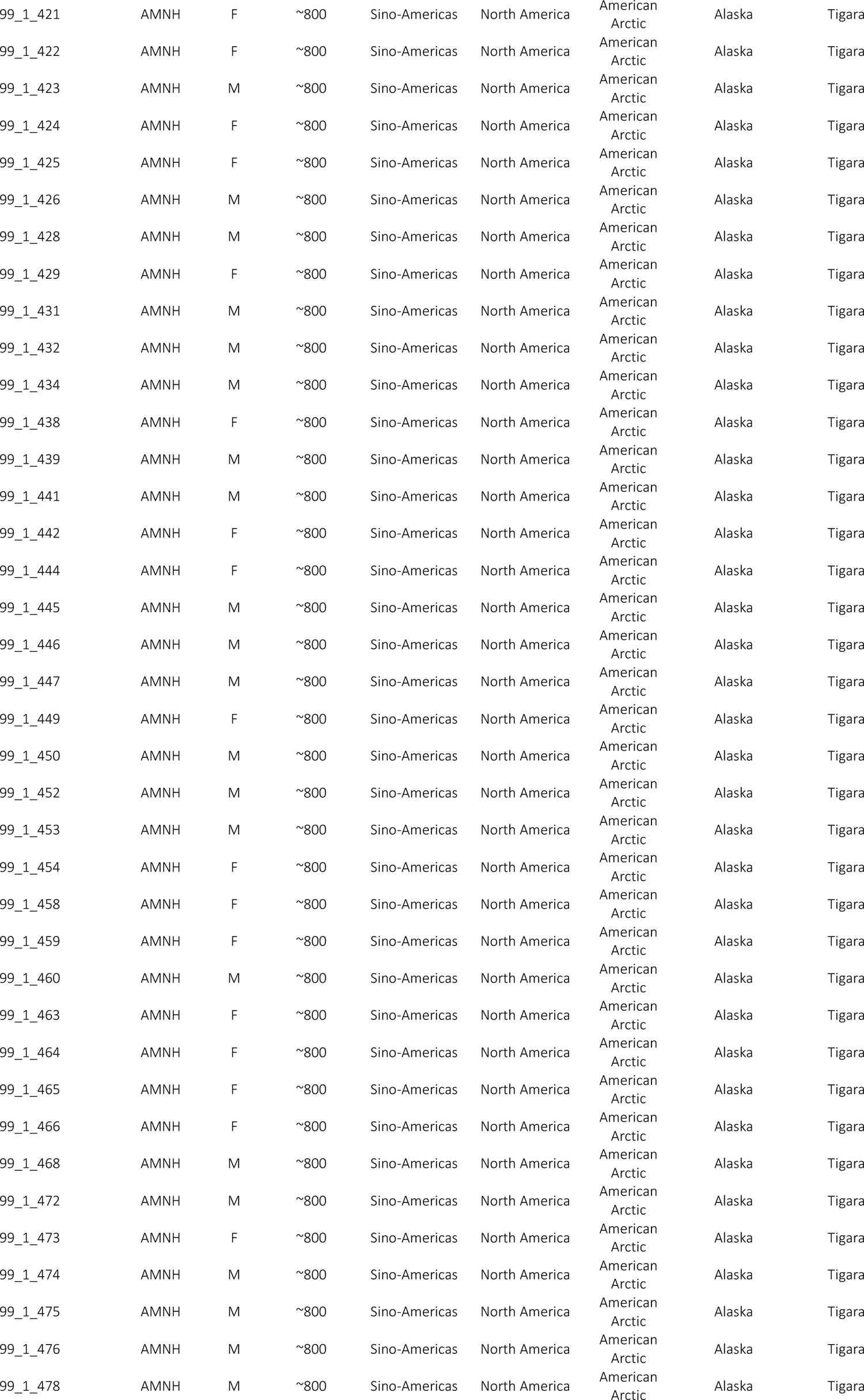
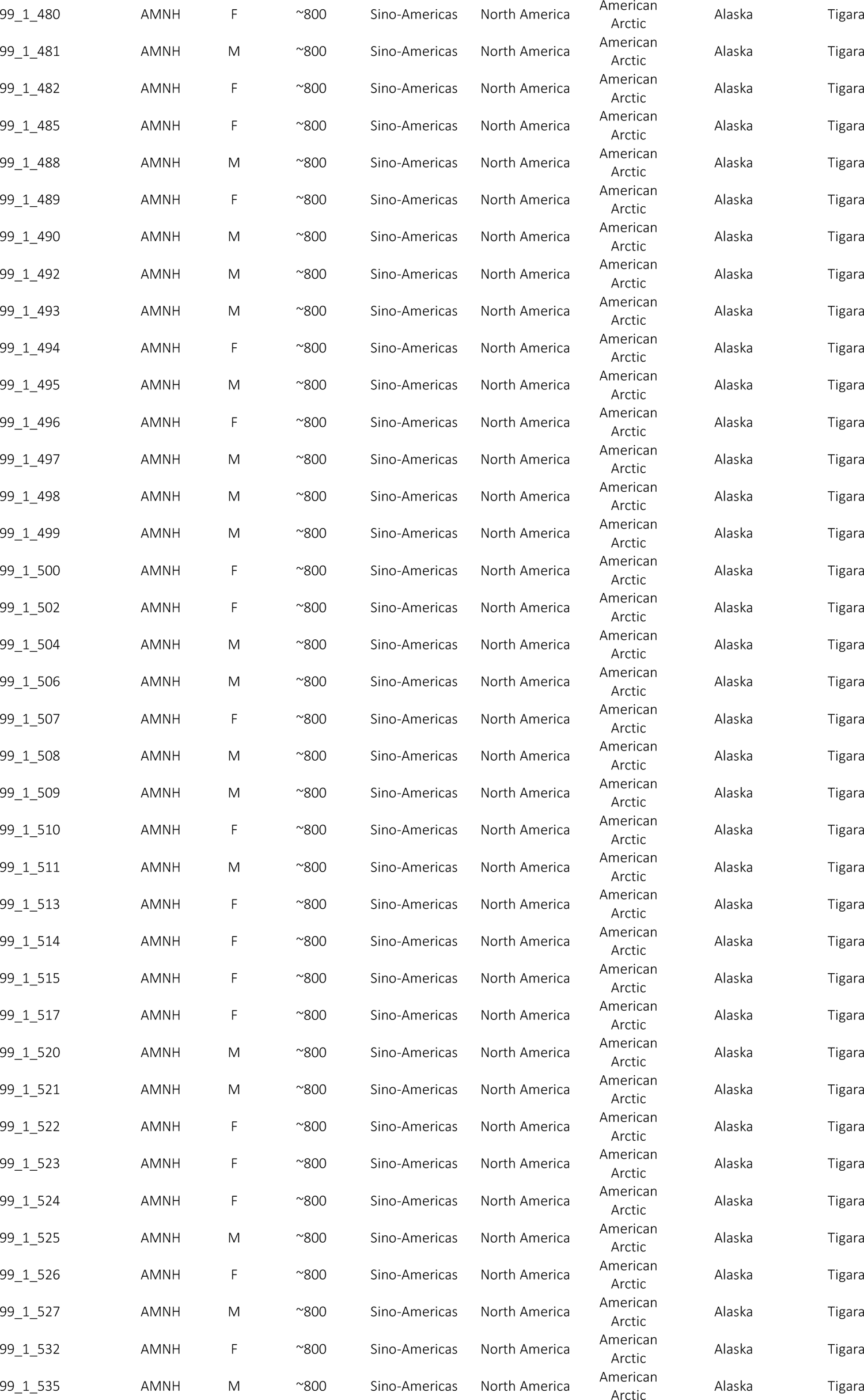
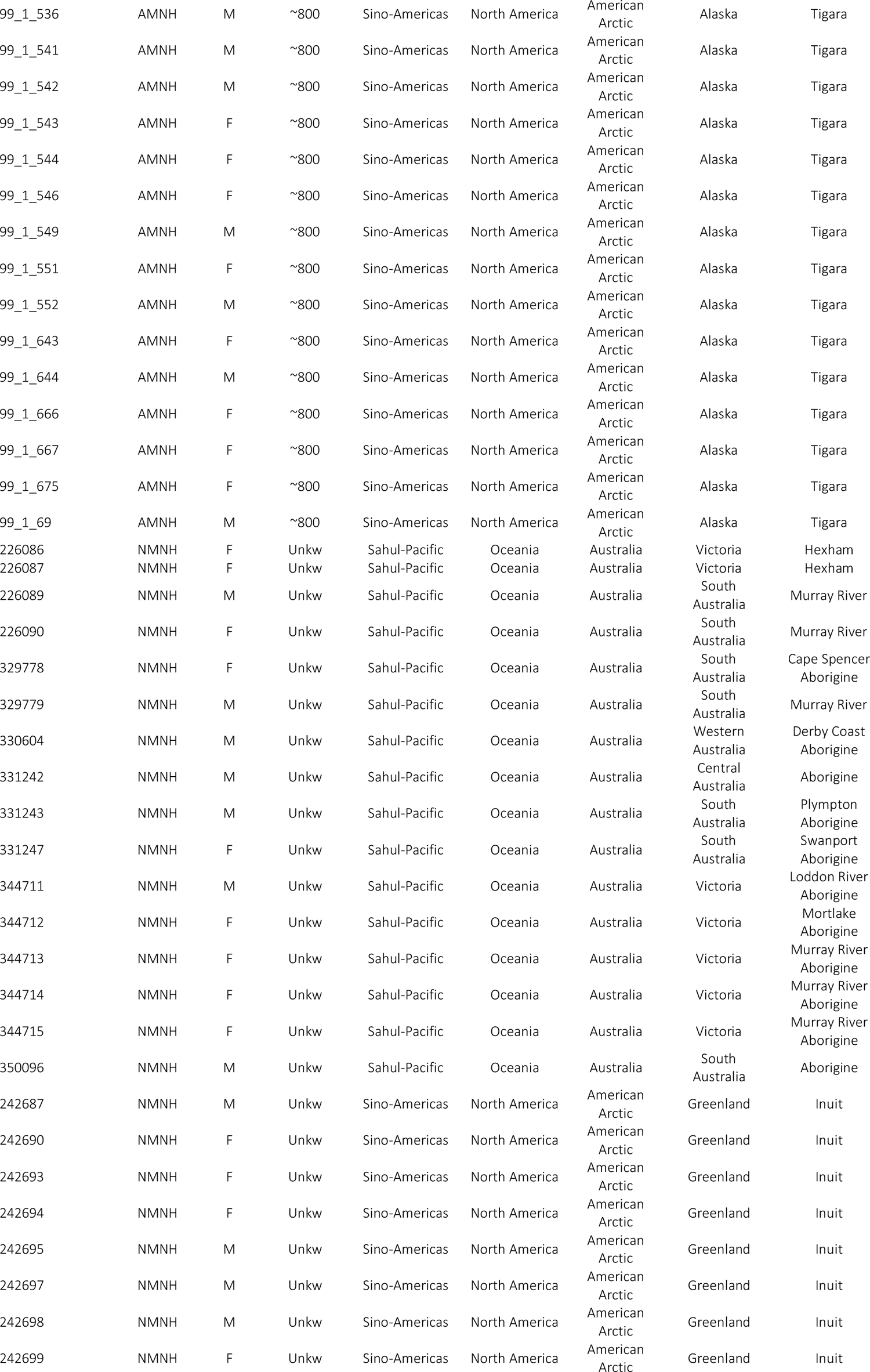
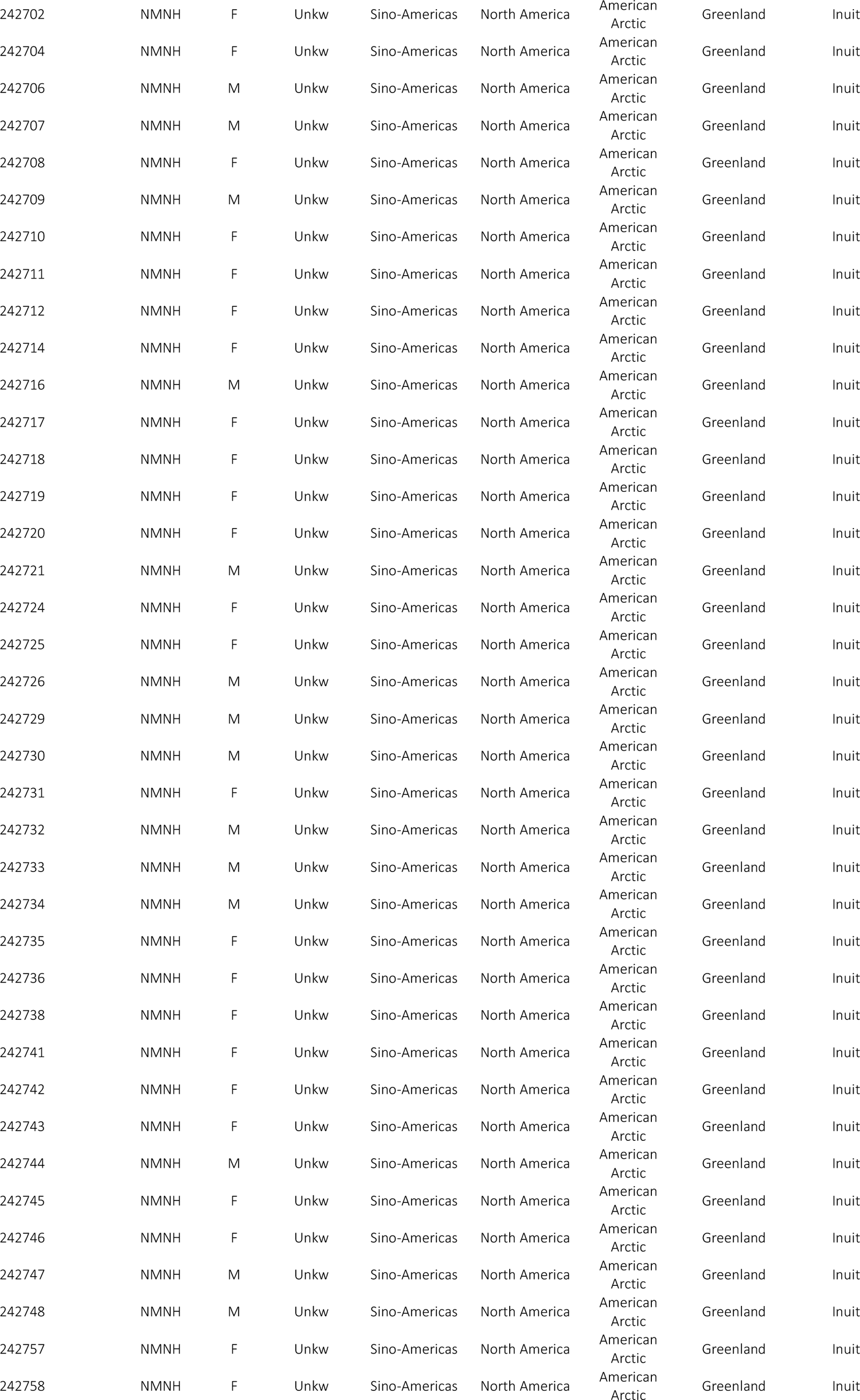
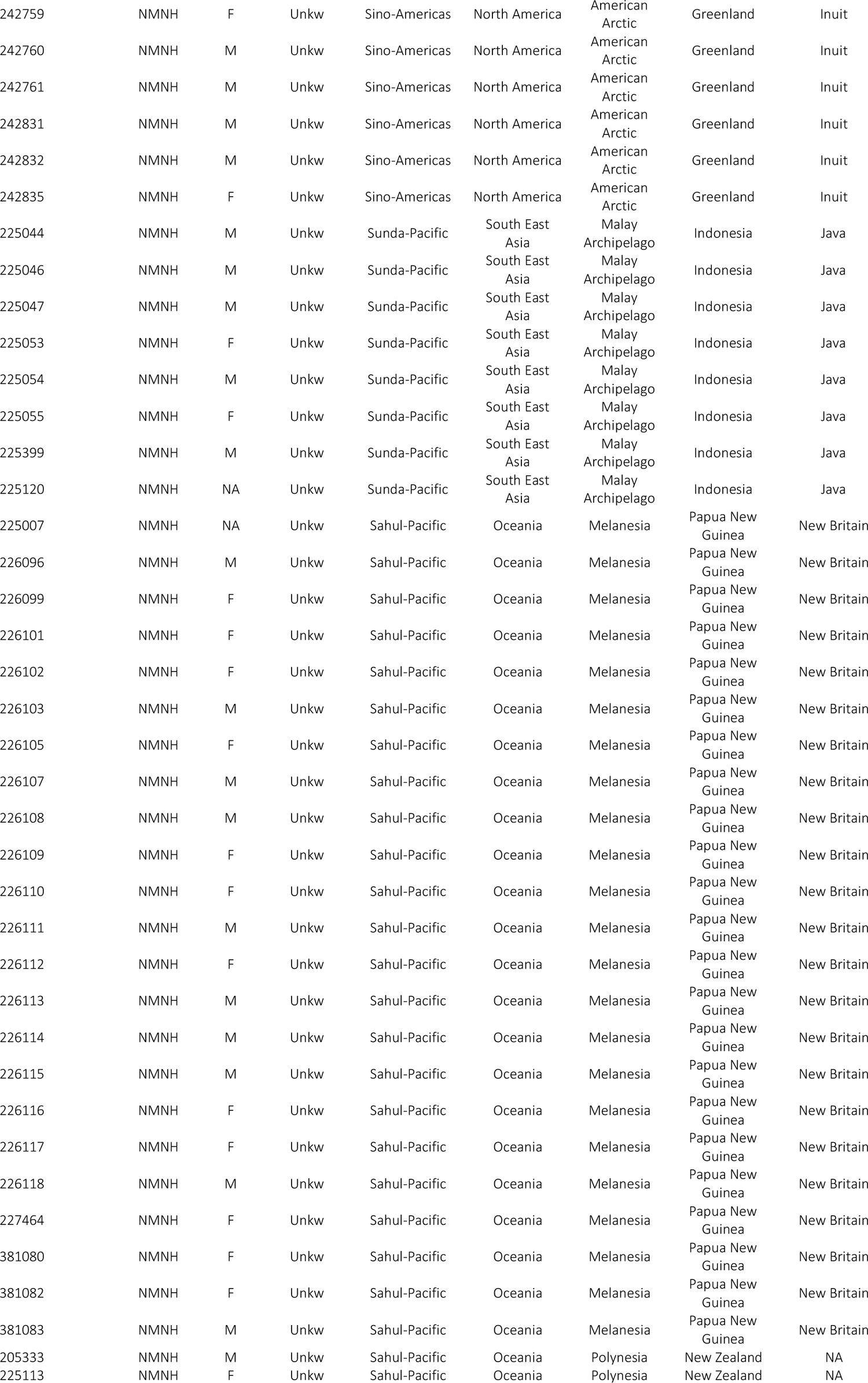
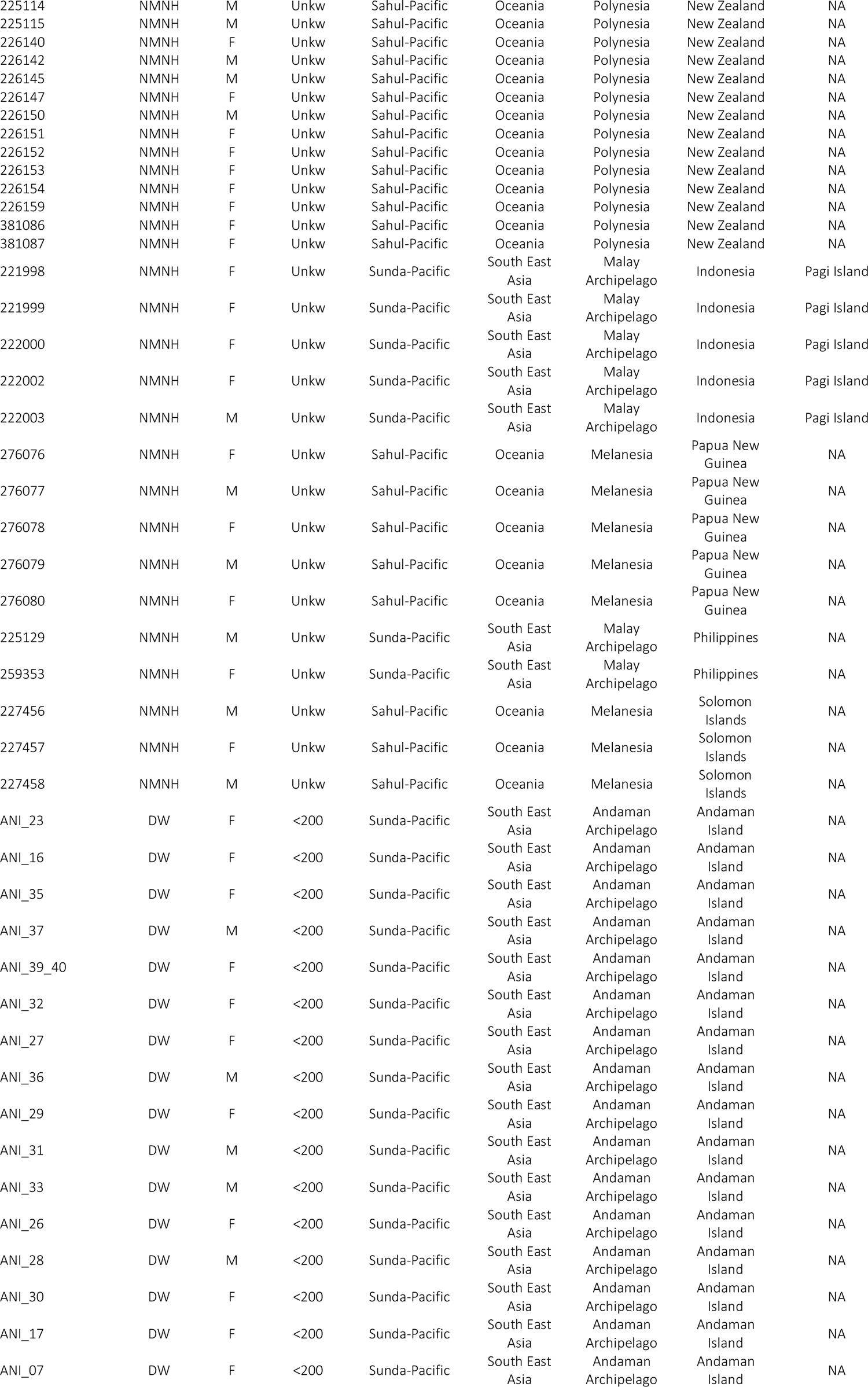
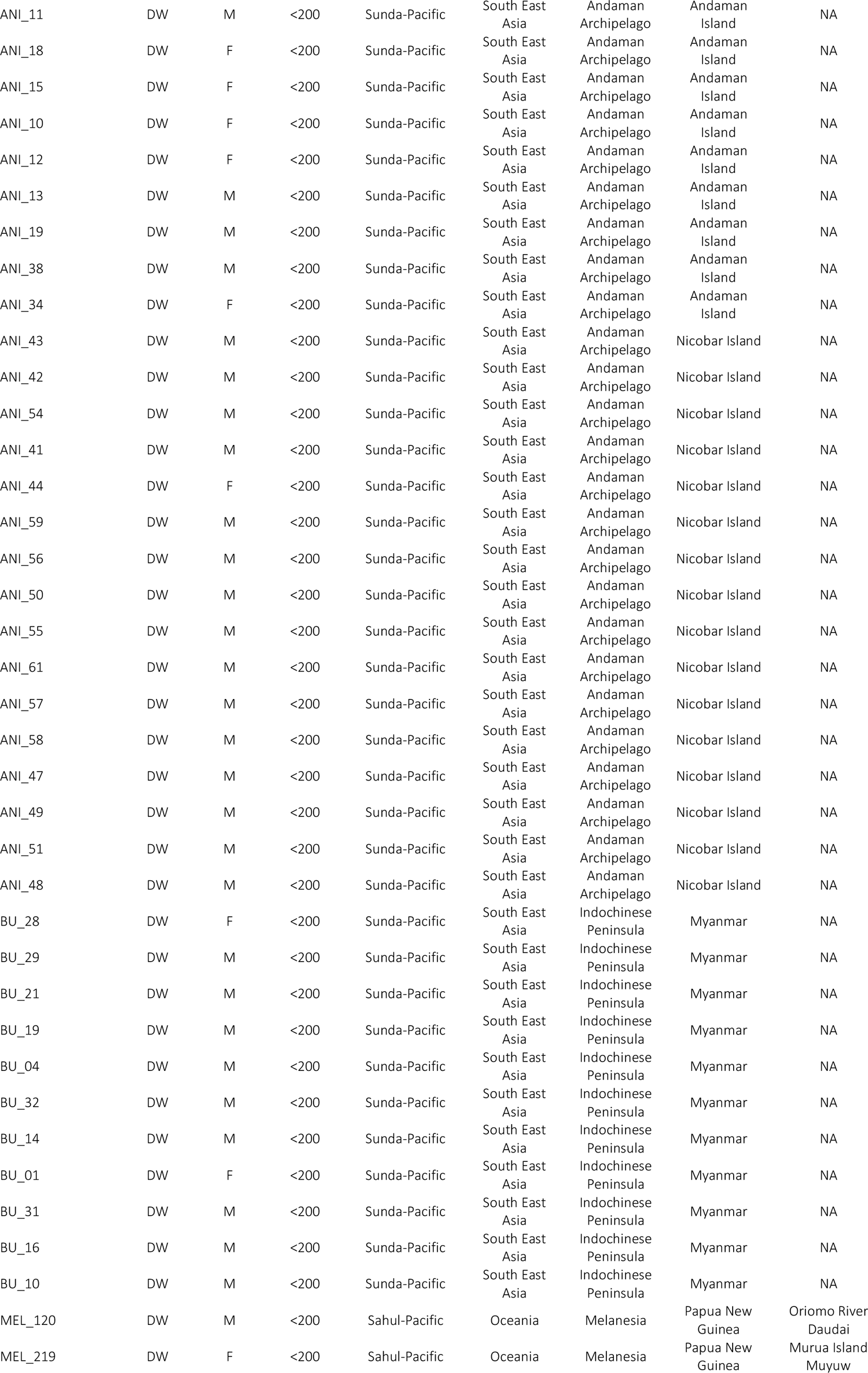
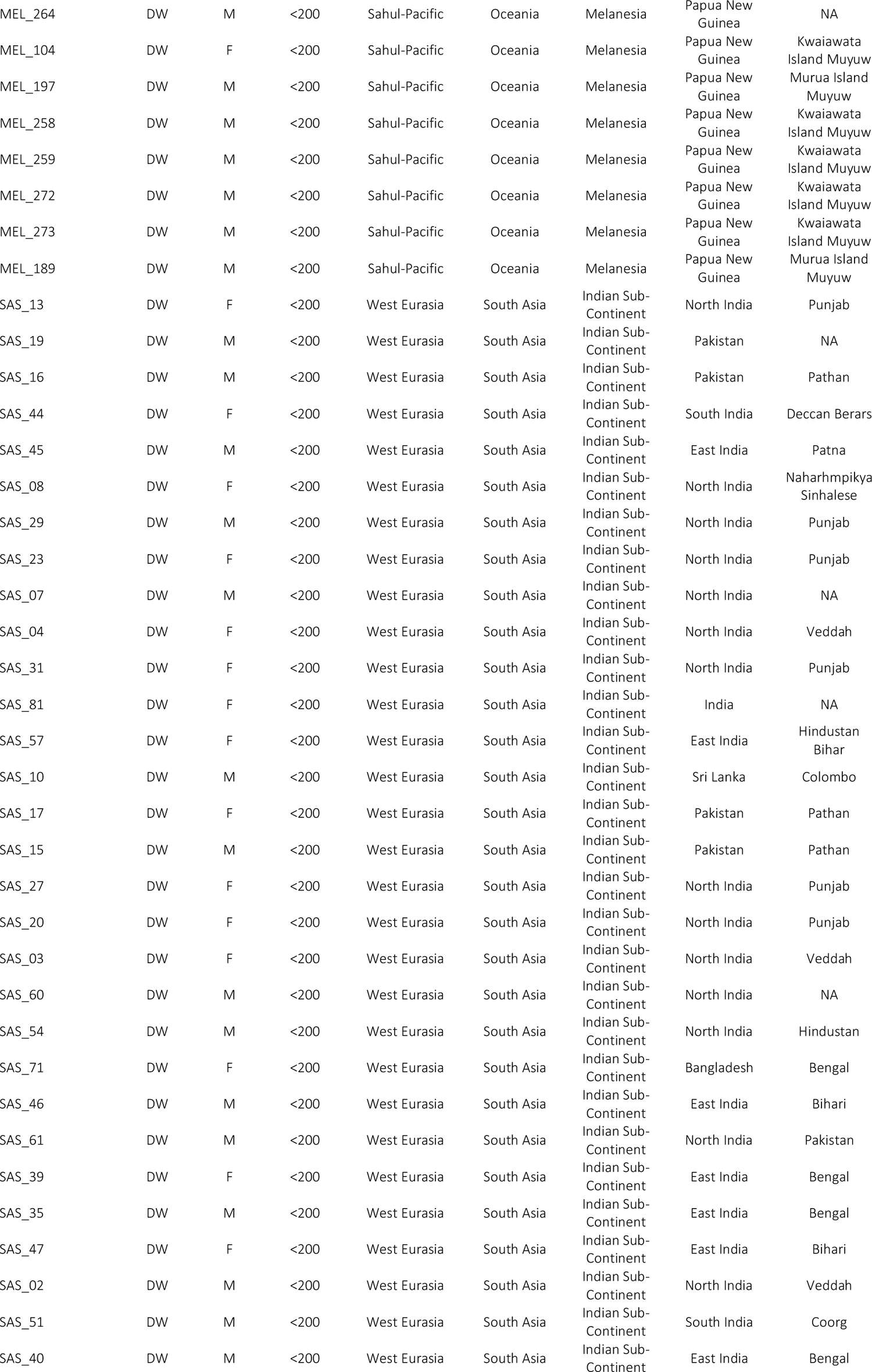
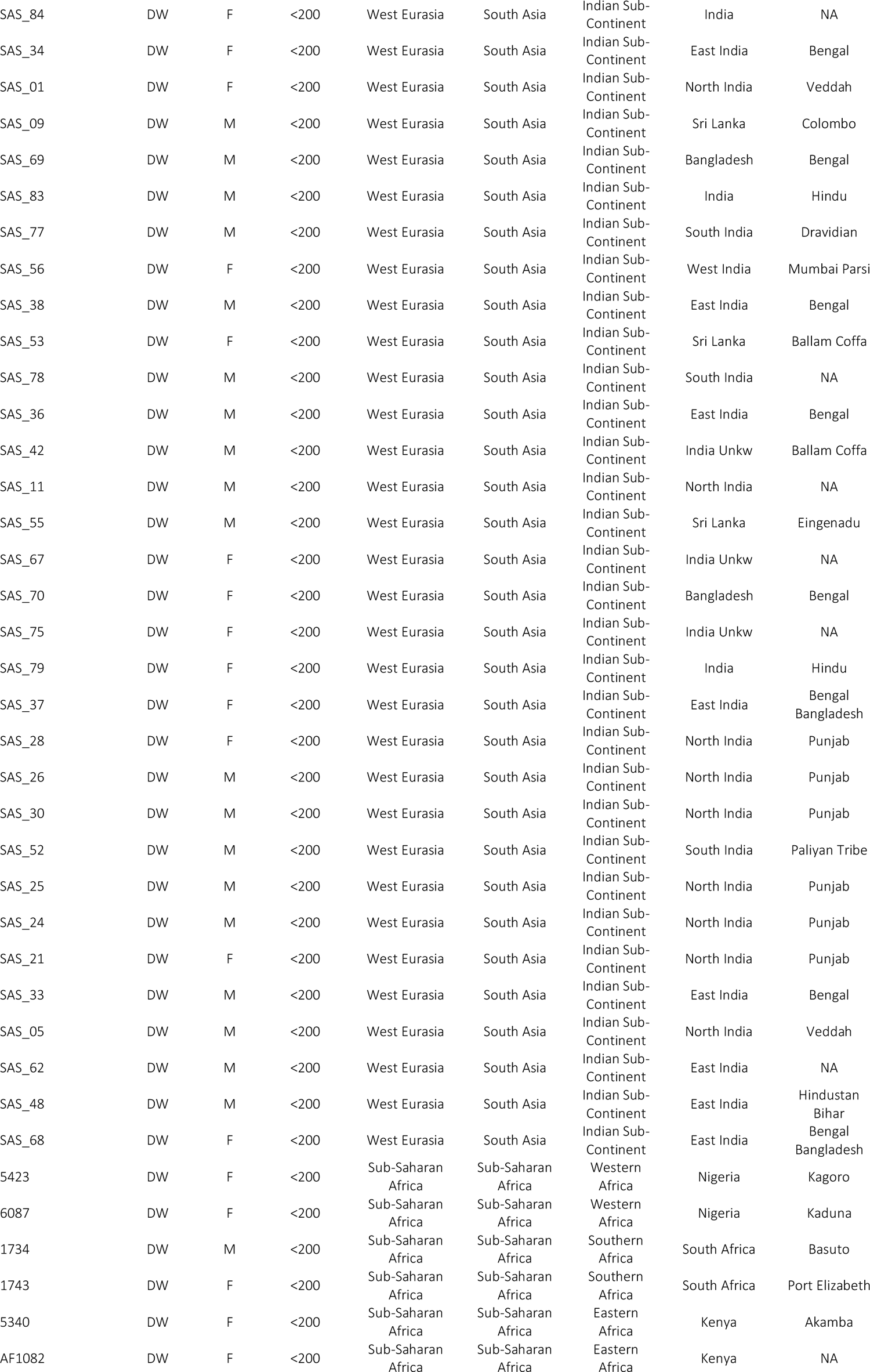
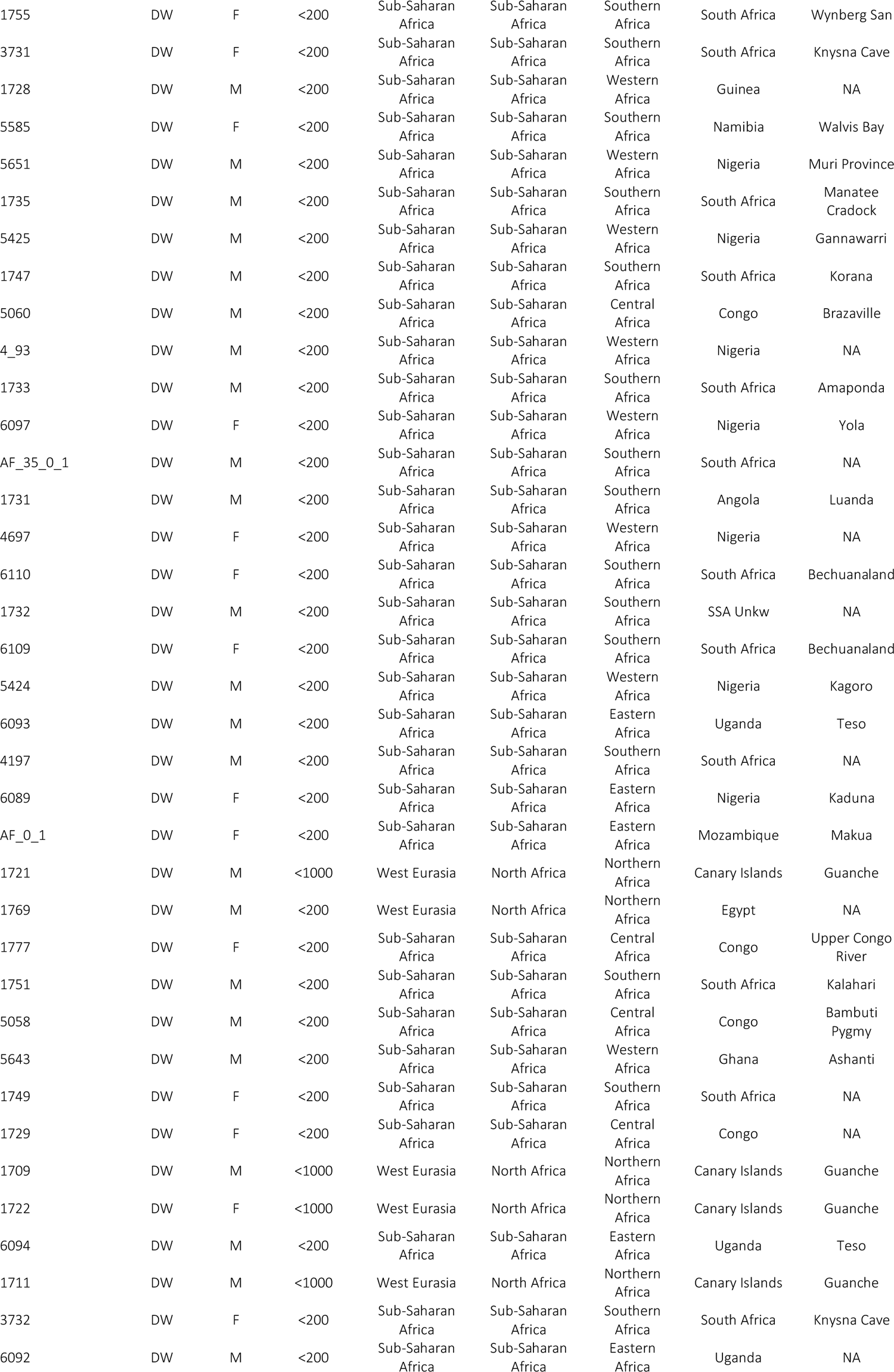
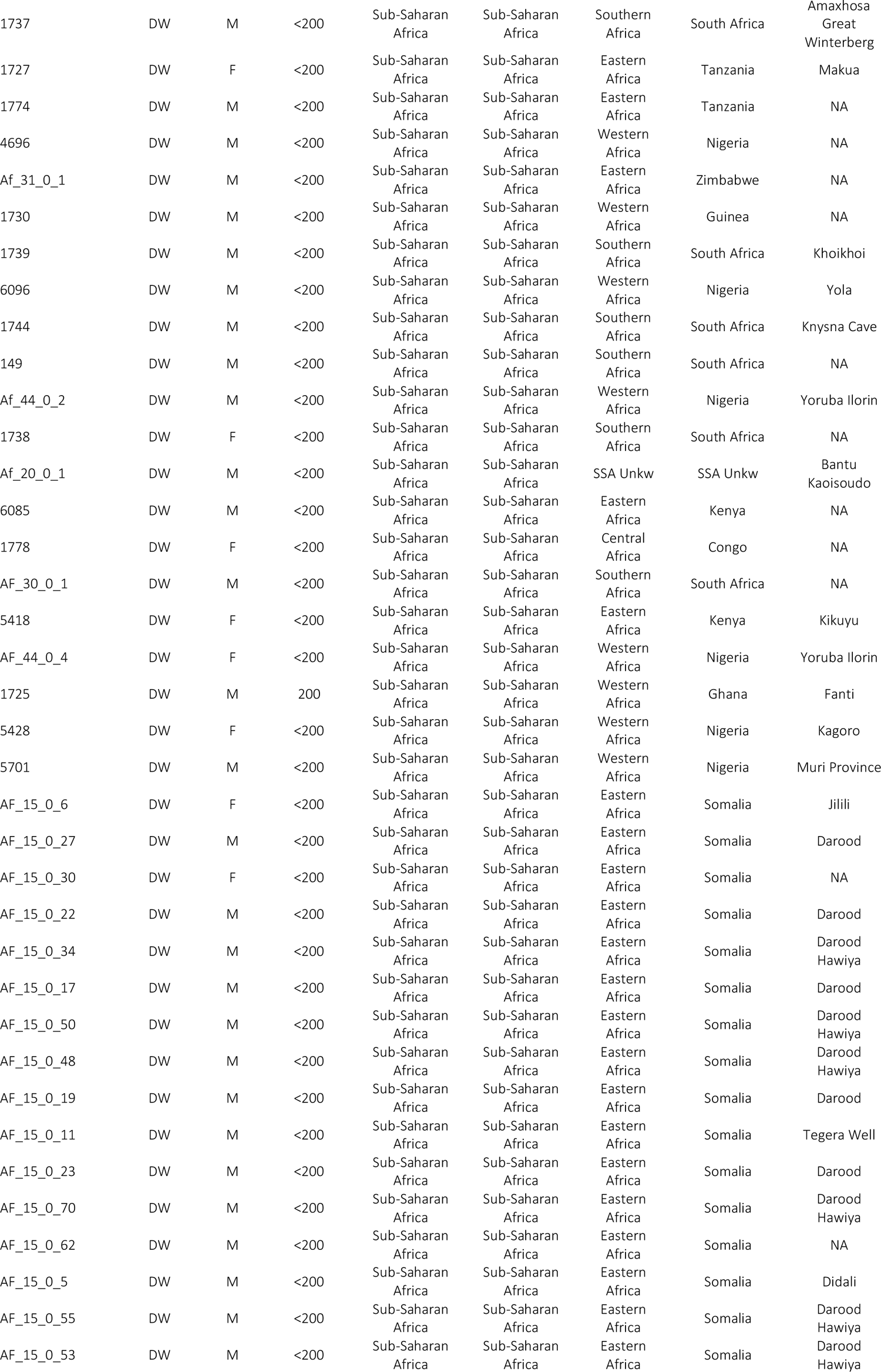
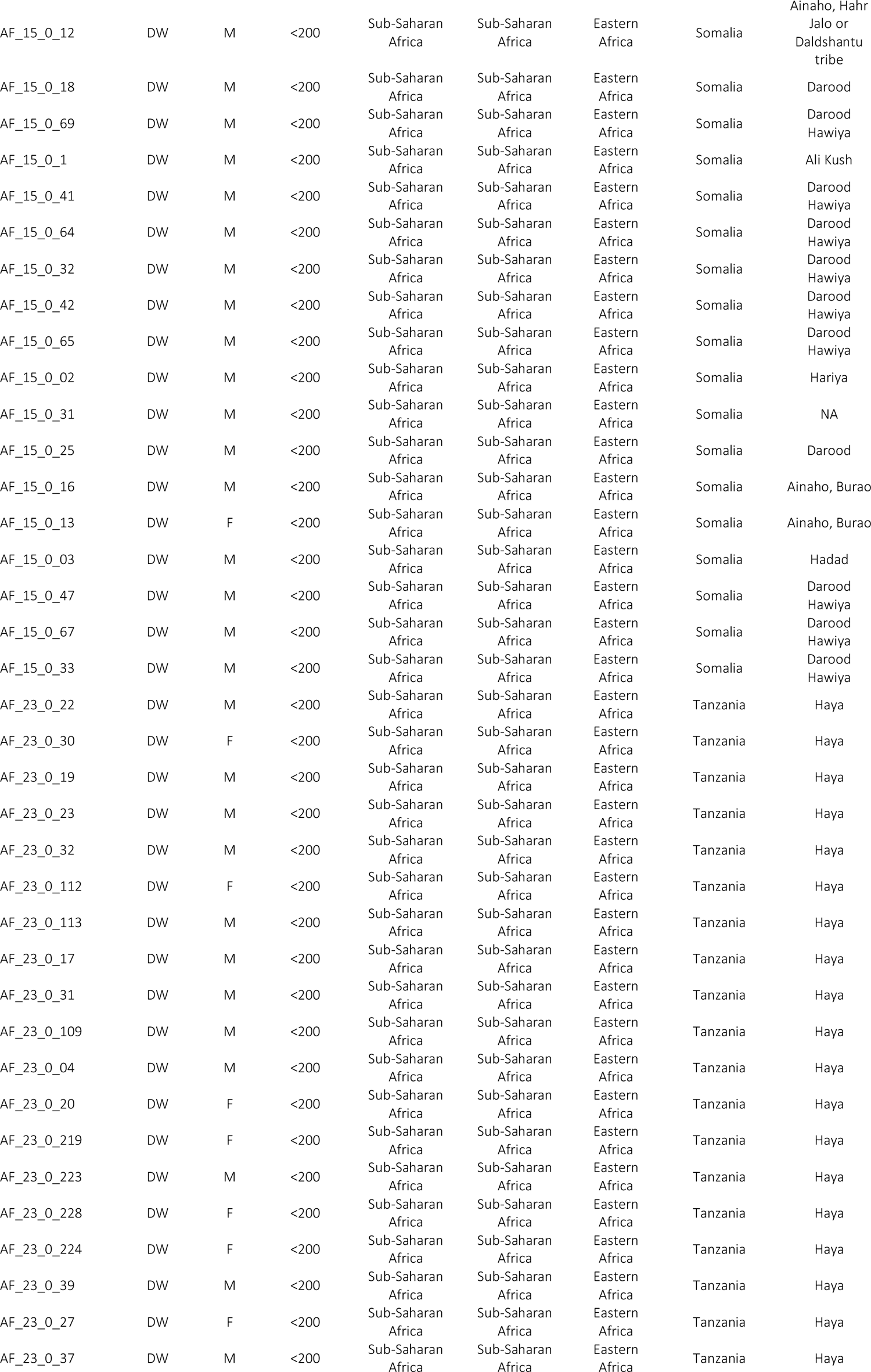
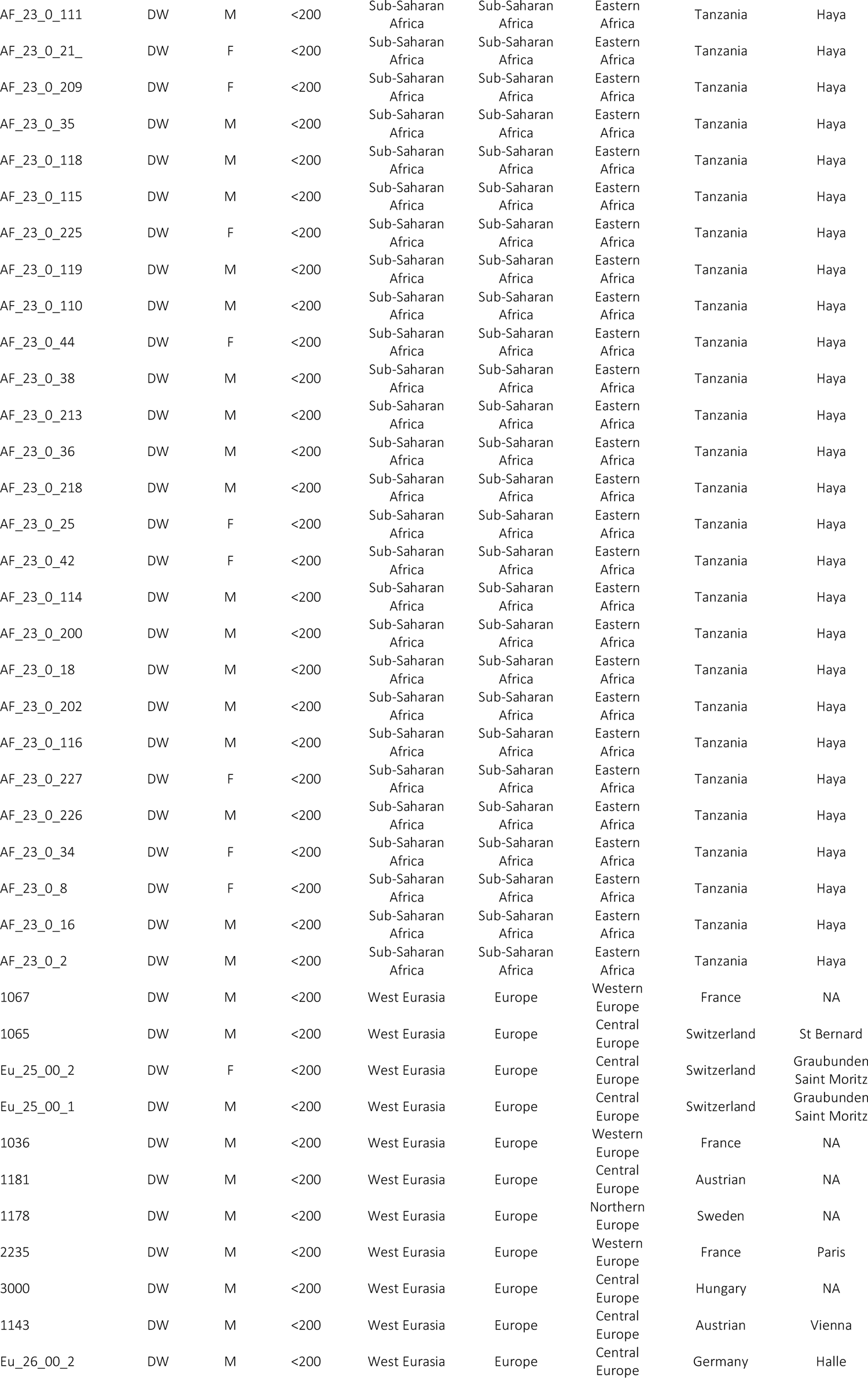
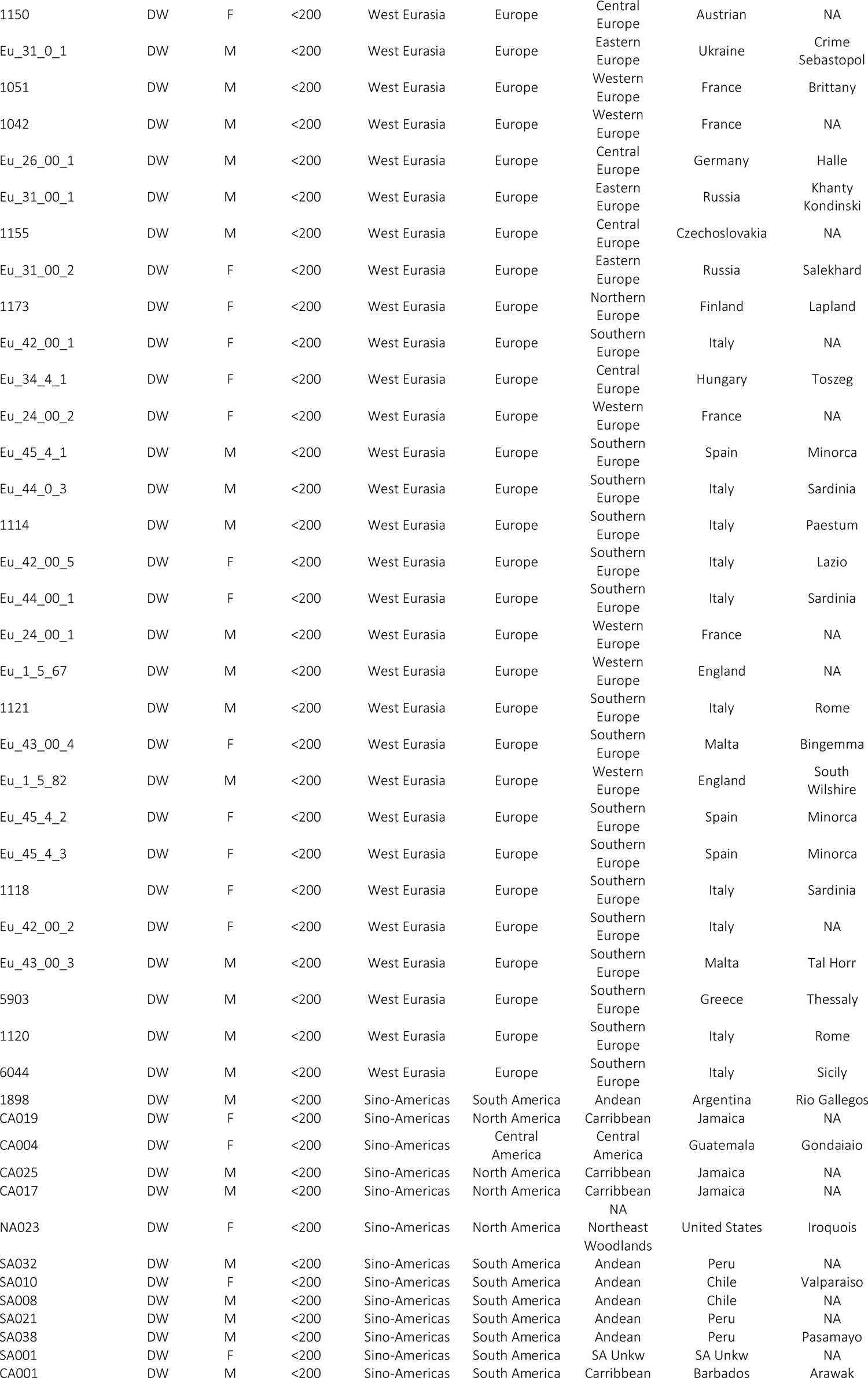
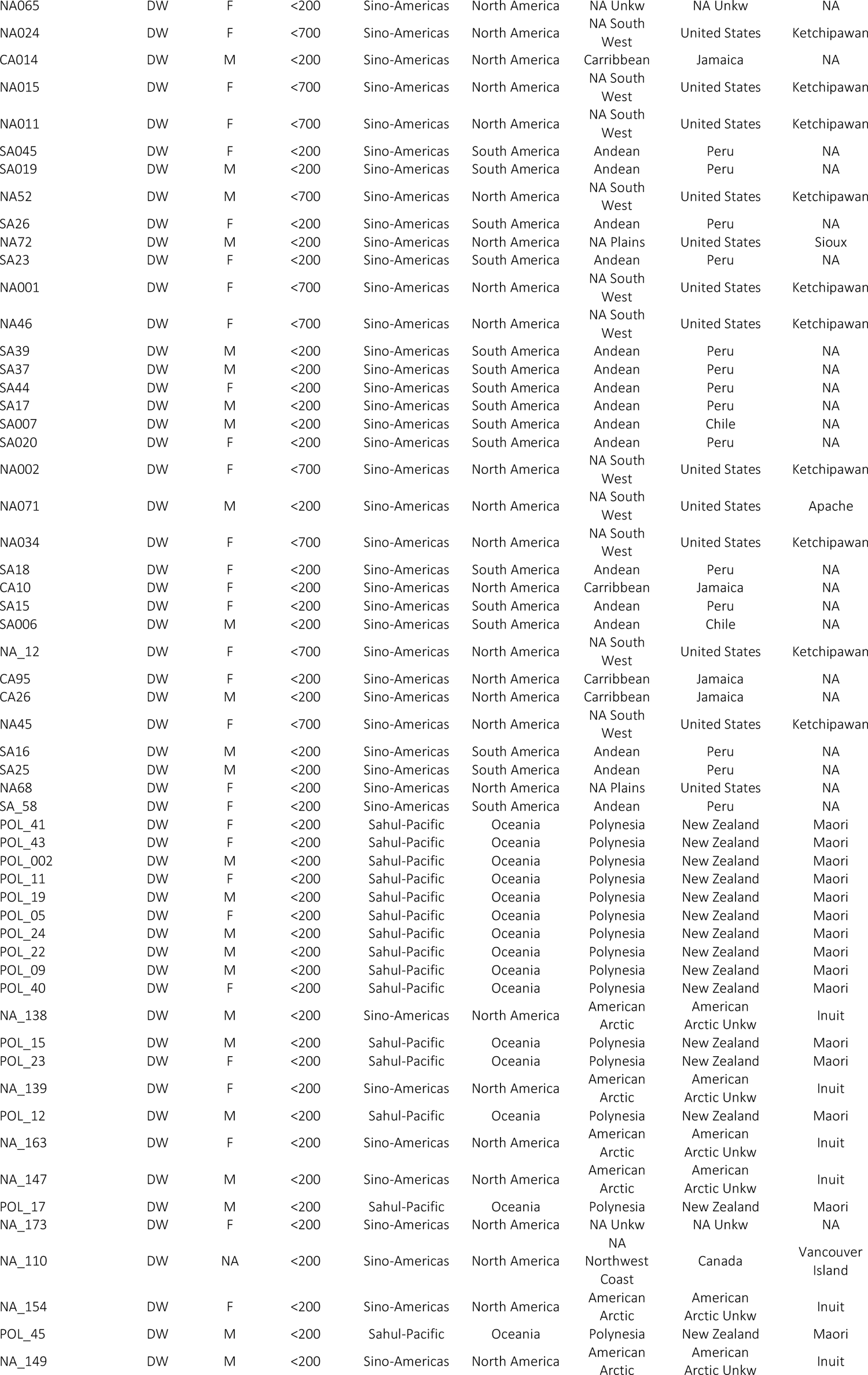
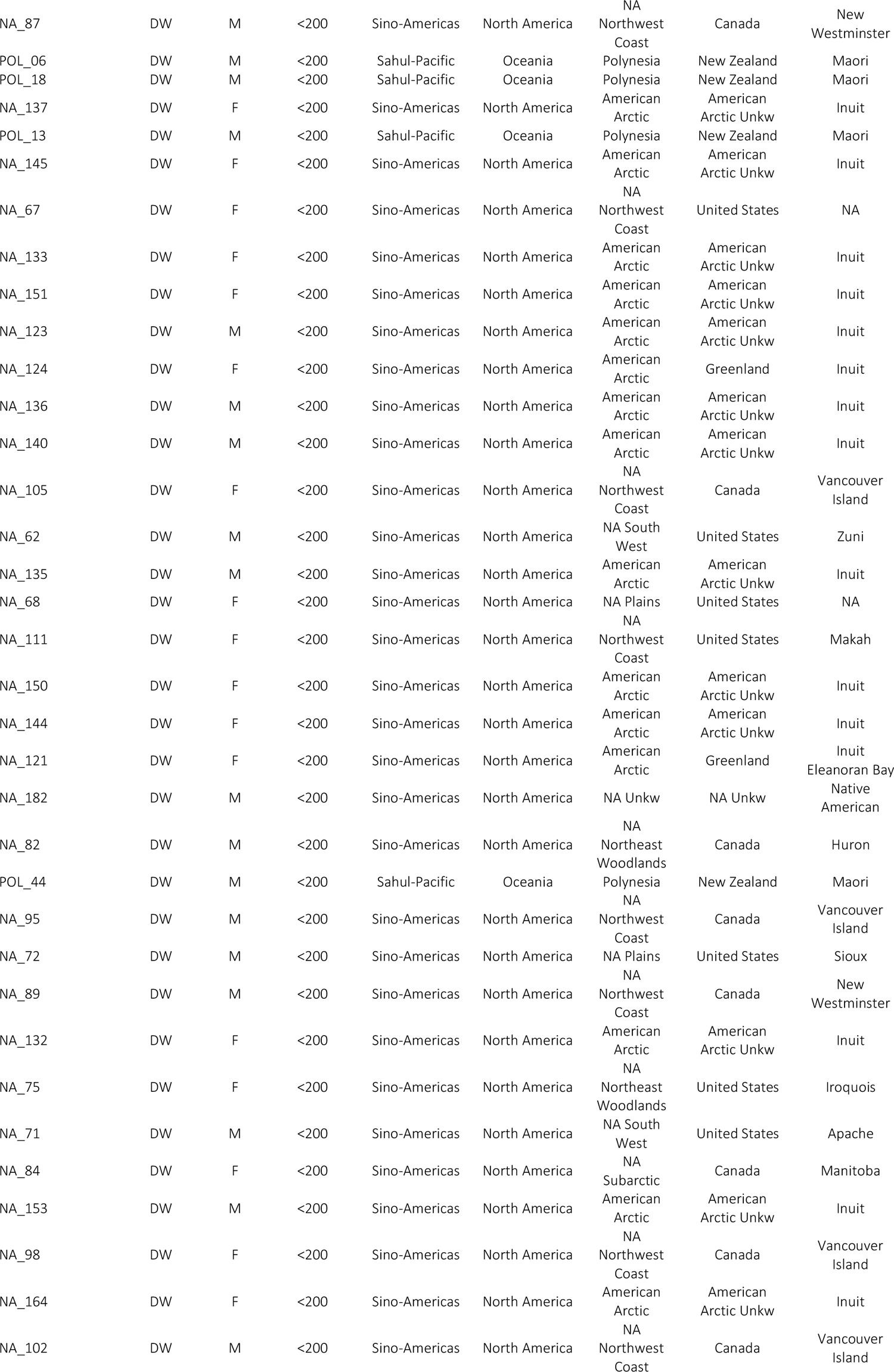
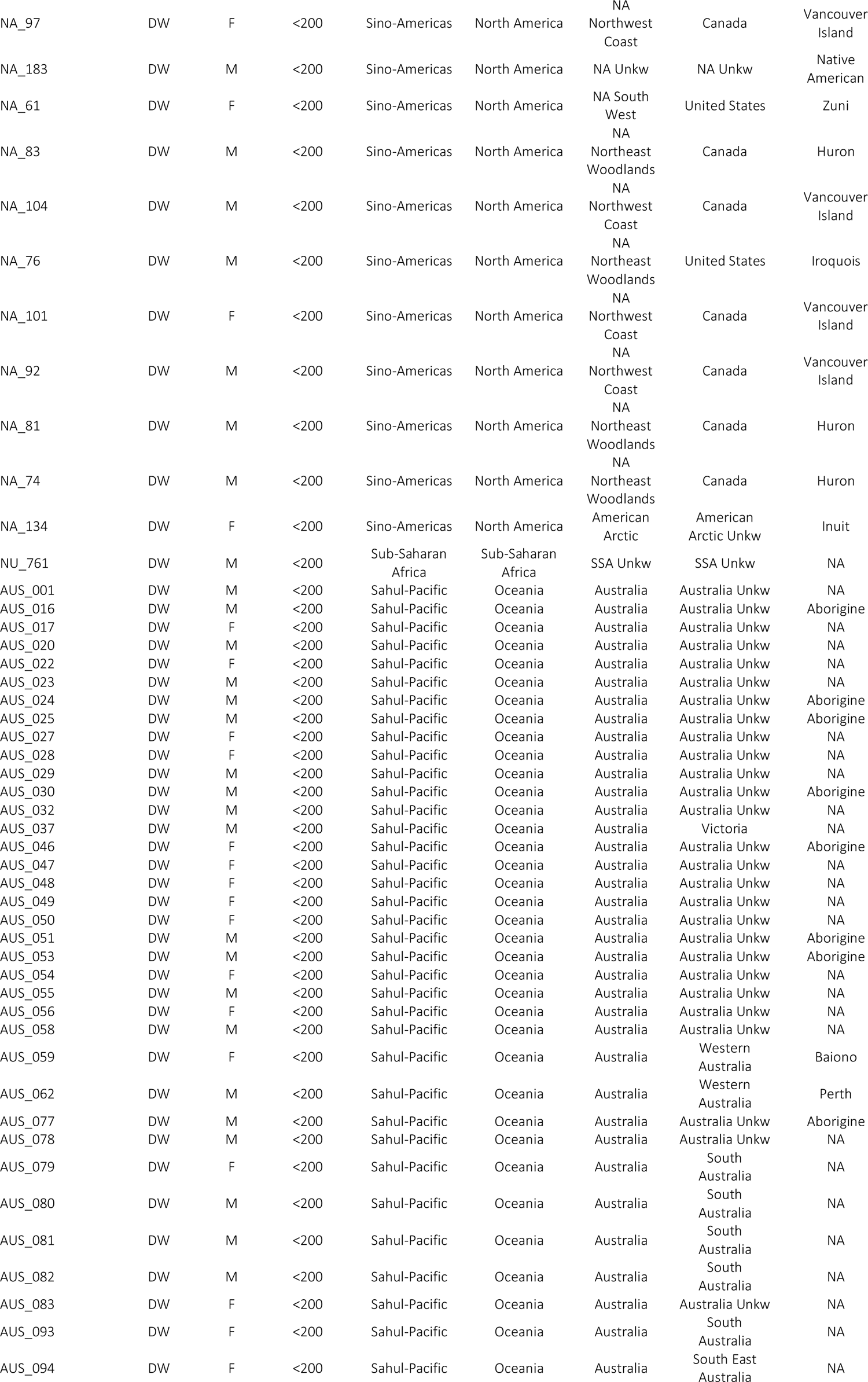
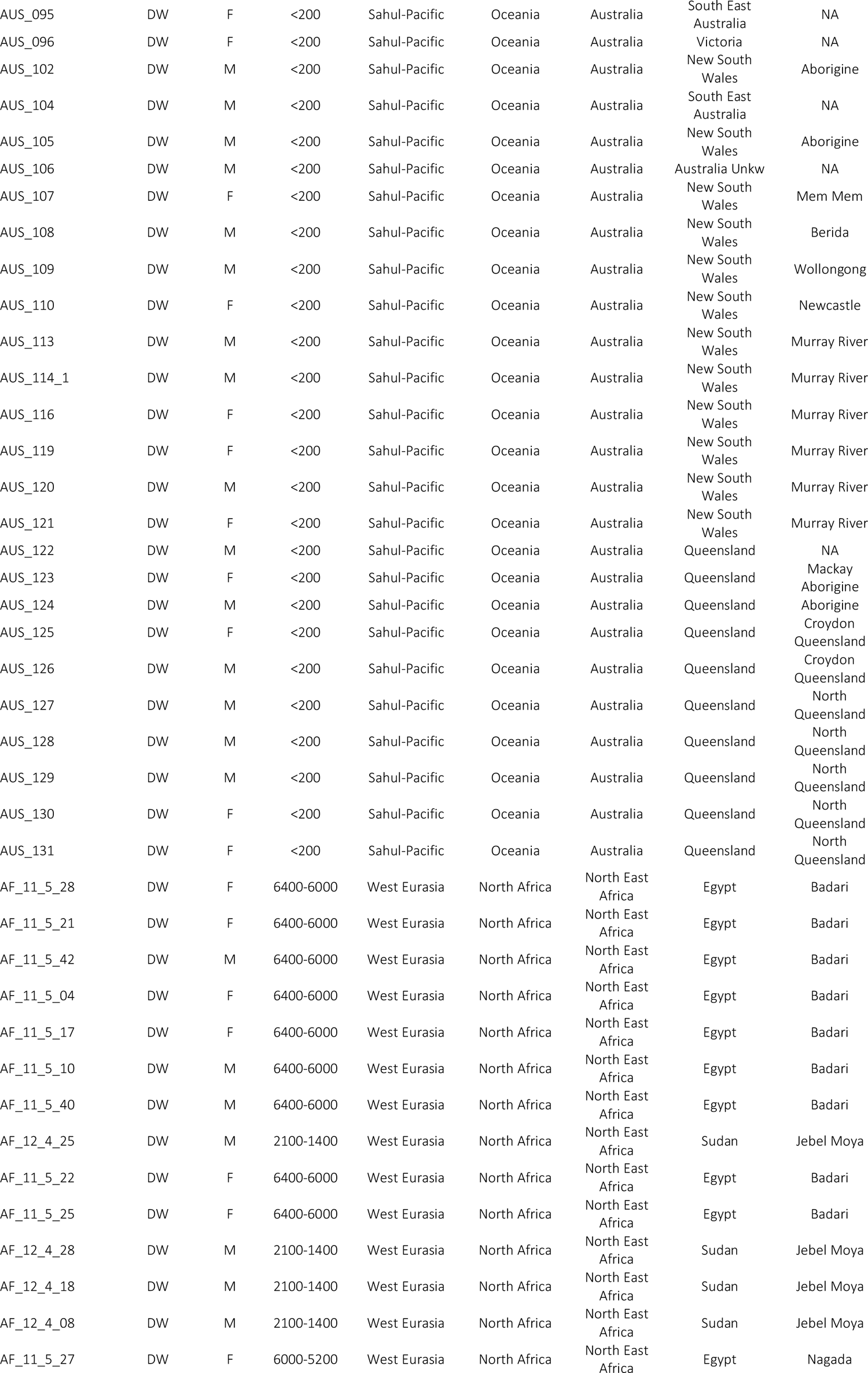
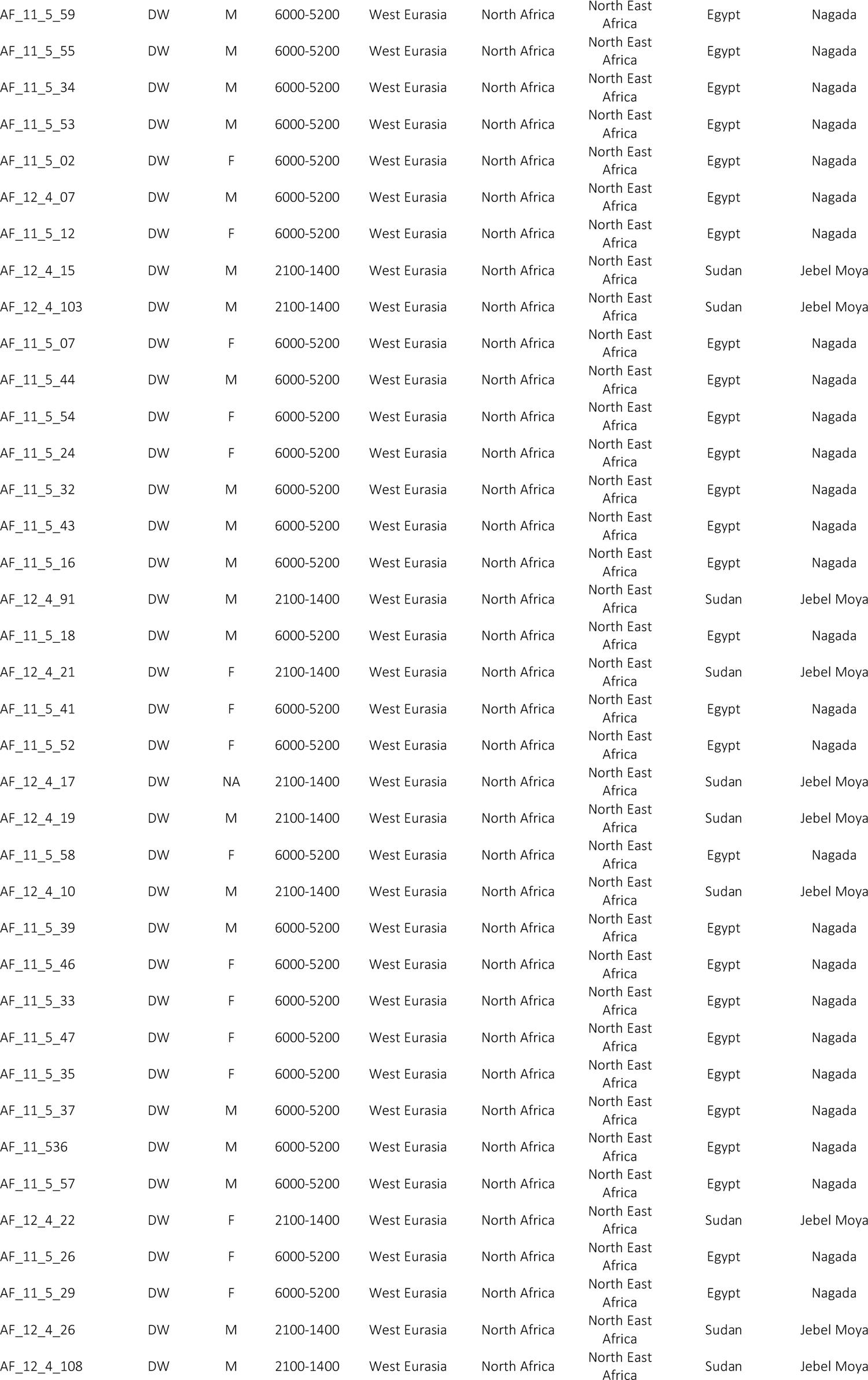
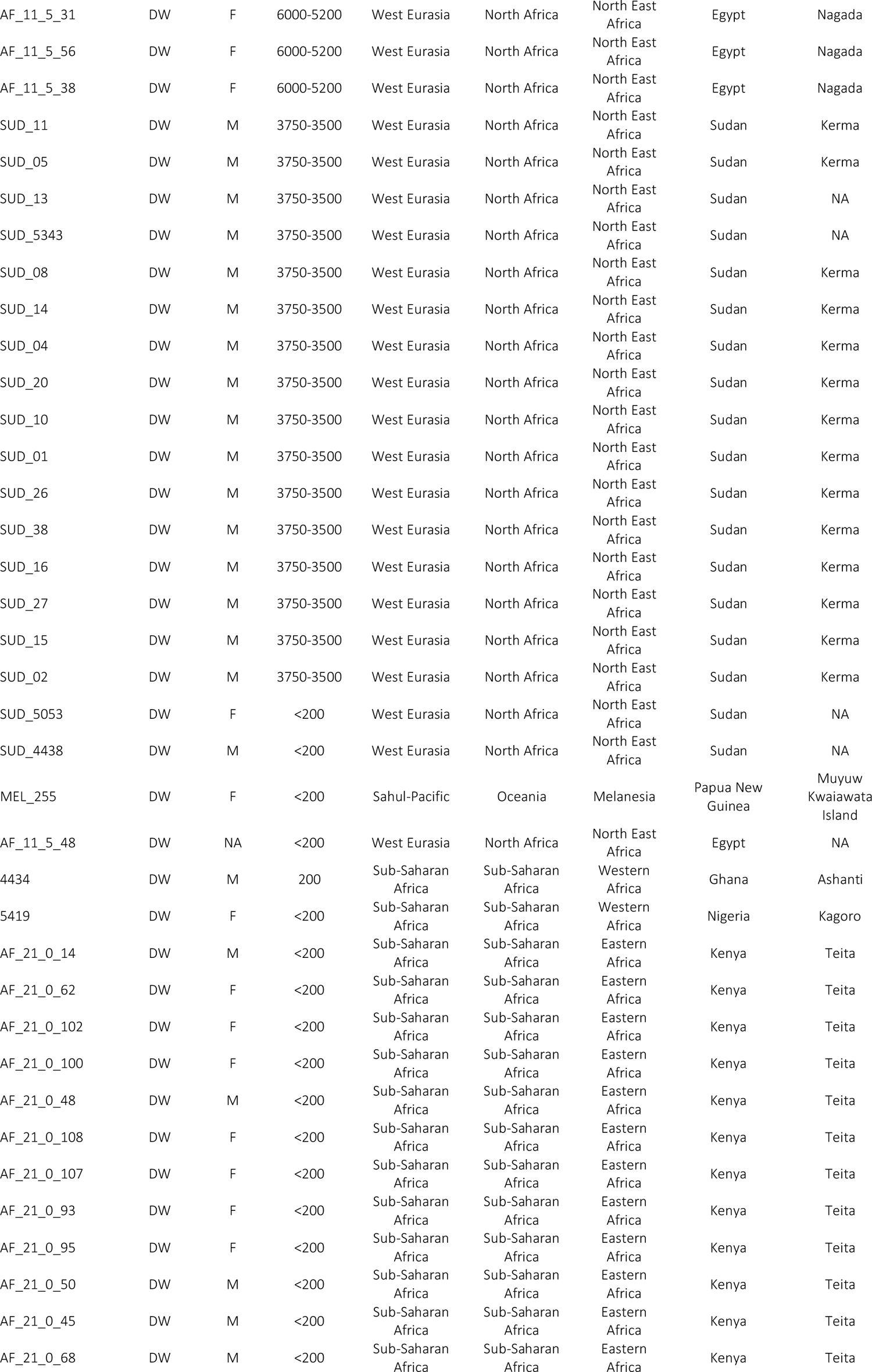
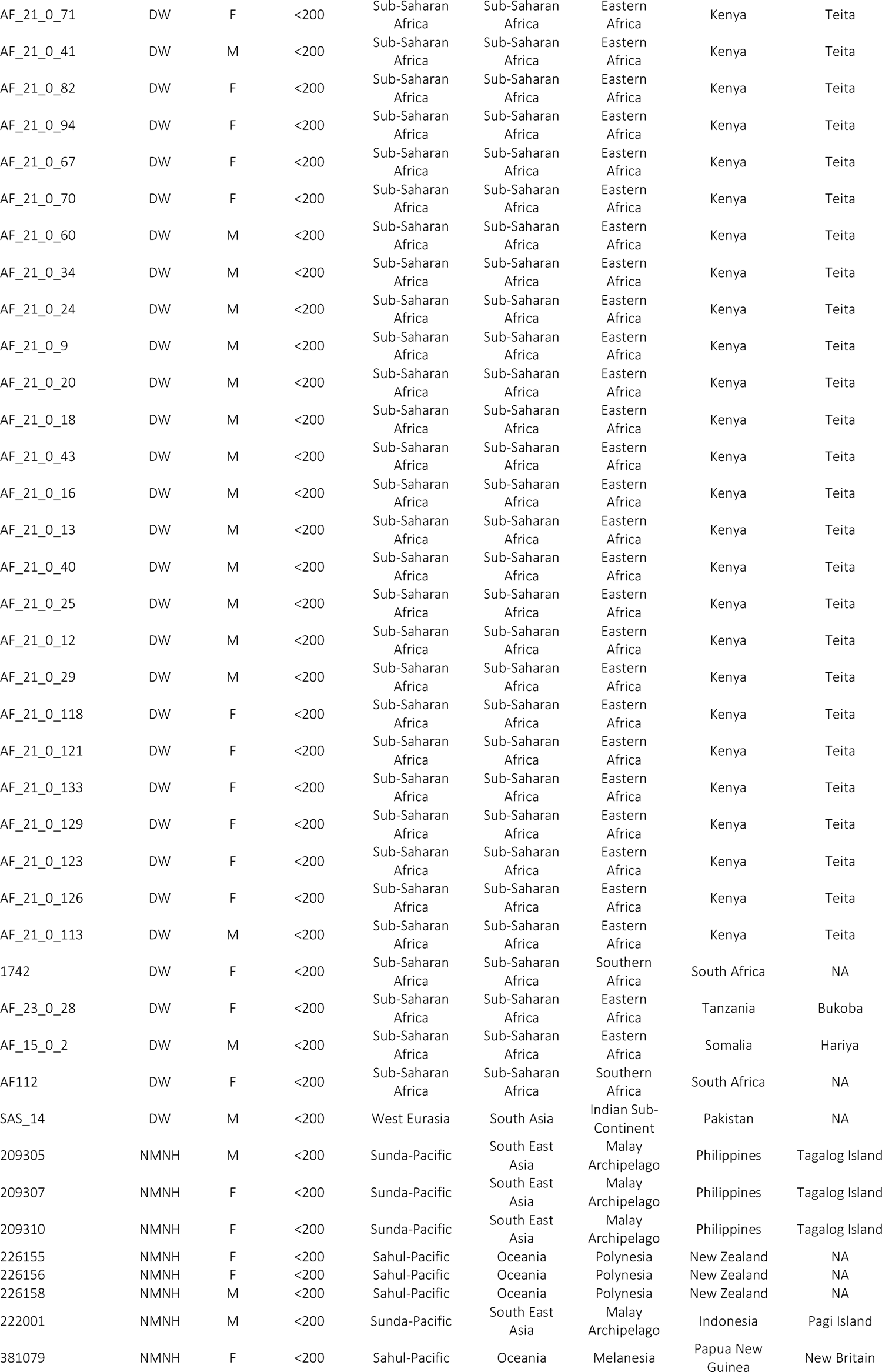
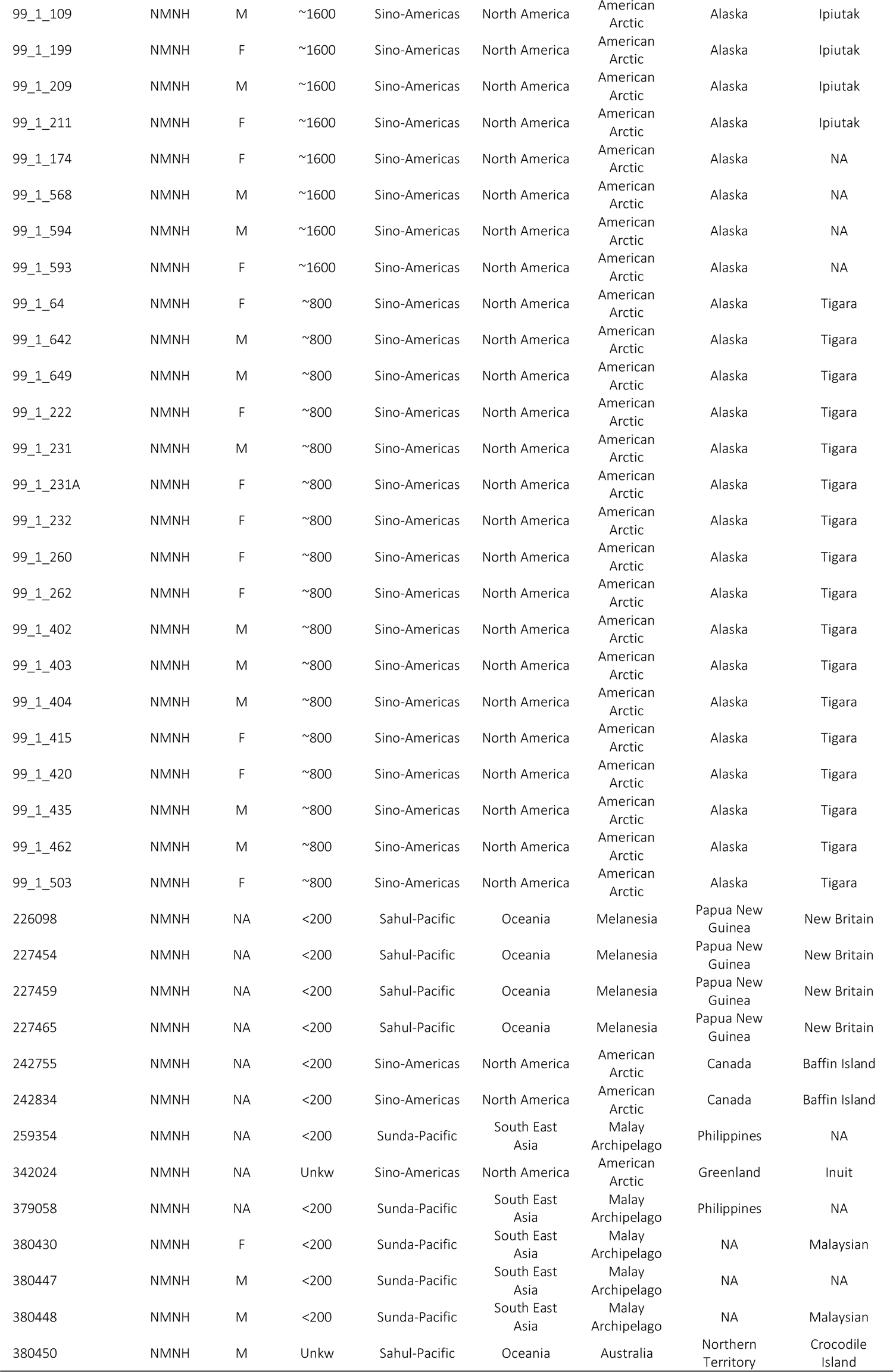
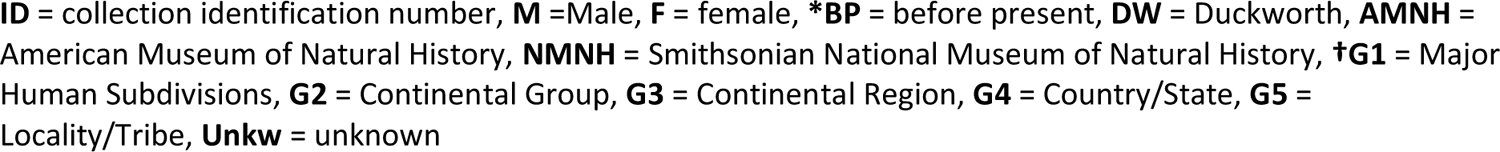
Collection information for individuals used in this study

**Table B:** Tukey Pair-wise comparisons of from PGLM model of canal to root number by tooth

| Contrast | Estimate | Odds Ratio | Std. Error | z-value | P-value |
| --- | --- | --- | --- | --- | --- |
| P<sup>3</sup> - P<sup>4</sup> | 0.0911 | 1.0953 | 0.0123 | 7.4180 | <.00001 |
| P<sup>3</sup> - M<sup>1</sup> | -0.6923 | 0.5004 | 0.0165 | -41.9106 | <.00001 |
| P<sup>3</sup> - M<sup>2</sup> | -0.6476 | 0.5233 | 0.0165 | -39.2830 | <.00001 |
| P<sup>3</sup> - M<sup>3</sup> | -0.6500 | 0.5221 | 0.0164 | -39.6955 | <.00001 |
| P<sup>3</sup> - P<sub>3</sub> | 0.1272 | 1.1356 | 0.0129 | 9.8777 | <.00001 |
| P<sup>3</sup> - P<sub>4</sub> | 0.1197 | 1.1272 | 0.0128 | 9.3303 | <.00001 |
| P<sup>3</sup> - M<sub>1</sub> | -0.2871 | 0.7504 | 0.0186 | -15.4013 | <.00001 |
| P<sup>3</sup> - M<sub>2</sub> | -0.3234 | 0.7237 | 0.0145 | -22.2508 | <.00001 |
| P<sup>3</sup> - M<sub>3</sub> | -0.3557 | 0.7006 | 0.0141 | -25.1822 | <.00001 |
| P<sup>4</sup> - M<sup>1</sup> | -0.7834 | 0.4569 | 0.0183 | -42.9011 | <.00001 |
| P<sup>4</sup> - M<sup>2</sup> | -0.7386 | 0.4778 | 0.0183 | -40.3220 | <.00001 |
| P<sup>4</sup> - M<sup>3</sup> | -0.7410 | 0.4766 | 0.0179 | -41.3743 | <.00001 |
| P<sup>4</sup> - P<sub>3</sub> | 0.0361 | 1.0368 | 0.0073 | 4.9477 | <.00001 |
| P<sup>4</sup> - P<sub>4</sub> | 0.0286 | 1.0290 | 0.0071 | 4.0477 | <.00001 |
| P<sup>4</sup> - M<sub>1</sub> | -0.3782 | 0.6851 | 0.0208 | -18.1390 | <.00001 |
| P<sup>4</sup> - M<sub>2</sub> | -0.4145 | 0.6607 | 0.0158 | -26.2447 | <.00001 |
| P<sup>4</sup> - M<sub>3</sub> | -0.4468 | 0.6397 | 0.0152 | -29.4156 | <.00001 |
| M<sup>1</sup> - M<sup>2</sup> | 0.0448 | 1.0458 | 0.0061 | 7.2915 | <.00001 |
| M<sup>1</sup> - M<sup>3</sup> | 0.0423 | 1.0433 | 0.0072 | 5.8984 | 0.0001 |
| M<sup>1</sup> - P<sub>3</sub> | 0.8195 | 2.2694 | 0.0174 | 47.1051 | <.00001 |
| M<sup>1</sup> - P<sub>4</sub> | 0.8120 | 2.2525 | 0.0178 | 45.6536 | **<.00001** |
| M<sup>1</sup> - M<sub>1</sub> | 0.4052 | 1.4997 | 0.0055 | 73.7824 | **<.00001** |
| M<sup>1</sup> - M<sub>2</sub> | 0.3689 | 1.4461 | 0.0053 | 69.3361 | **<.00001** |
| M<sup>1</sup> - M<sub>3</sub> | 0.3366 | 1.4002 | 0.0055 | 61.0848 | **<.00001** |
| M<sup>2</sup> - M<sup>3</sup> | -0.0024 | 0.9976 | 0.0078 | -0.3101 | 0.0222 |
| M<sup>2</sup> - P<sub>3</sub> | 0.7748 | 2.1701 | 0.0177 | 43.6840 | **<.00001** |
| M<sup>2</sup> - P<sub>4</sub> | 0.7673 | 2.1539 | 0.0181 | 42.3303 | **<.00001** |
| M<sup>2</sup> - M<sub>1</sub> | 0.3605 | 1.4340 | 0.0085 | 42.4873 | **<.00001** |
| M<sup>2</sup> - M<sub>2</sub> | 0.3241 | 1.3828 | 0.0071 | 45.3929 | **<.00001** |
| M<sup>2</sup> - M<sub>3</sub> | 0.2918 | 1.3389 | 0.0074 | 39.2476 | **<.00001** |
| M<sup>3</sup> - P<sub>3</sub> | 0.7772 | 2.1753 | 0.0174 | 44.7211 | **<.00001** |
| M<sup>3</sup> - P<sub>4</sub> | 0.7697 | 2.1591 | 0.0177 | 43.4366 | **<.00001** |
| M<sup>3</sup> - M<sub>1</sub> | 0.3629 | 1.4375 | 0.0094 | 38.4709 | **<.00001** |
| M<sup>3</sup> - M<sub>2</sub> | 0.3265 | 1.3862 | 0.0078 | 41.6325 | **<.00001** |
| M<sup>3</sup> - M<sub>3</sub> | 0.2942 | 1.3421 | 0.0080 | 36.9683 | **<.00001**<sup>#</sup> |
| P<sub>3</sub> - P<sub>4</sub> | -0.0075 | 0.9925 | 0.0019 | -3.8619 | **<.00001** |
| P<sub>3</sub> - M<sub>1</sub> | -0.4143 | 0.6608 | 0.0200 | -20.6987 | **<.00001** |
| P<sub>3</sub> - M<sub>2</sub> | -0.4506 | 0.6372 | 0.0147 | -30.7287 | **<.00001** |
| P<sub>3</sub> - M<sub>3</sub> | -0.4829 | 0.6170 | 0.0141 | -34.2334 | **<.00001** |
| P<sub>4</sub> - M<sub>1</sub> | -0.4068 | 0.6658 | 0.0204 | -19.9481 | **<.00001** |
| P<sub>4</sub> - M<sub>2</sub> | -0.4431 | 0.6420 | 0.0151 | -29.3666 | **<.00001** |
| P<sub>4</sub> - M<sub>3</sub> | -0.4754 | 0.6216 | 0.0145 | -32.8780 | **<.00001** |
| M<sub>1</sub> - M<sub>2</sub> | -0.0364 | 0.9643 | 0.0078 | -4.6724 | **<.00001** |
| M<sub>1</sub> - M<sub>3</sub> | -0.0687 | 0.9336 | 0.0079 | -8.7160 | **<.00001** |
| M<sub>2</sub> - M<sub>3</sub> | -0.0323 | 0.9682 | 0.0050 | -6.4832 | **<.00001** |

<sup>#</sup> Does not meet Bonferroni correction of $0.05/45 \approx 0.001$ , where 45 is the number of pair-wise tests.\*
Significant values in bold after Bonferroni correction. Model fitted without intercept. Results are given on the log (not the response) scale.

**Table C:**
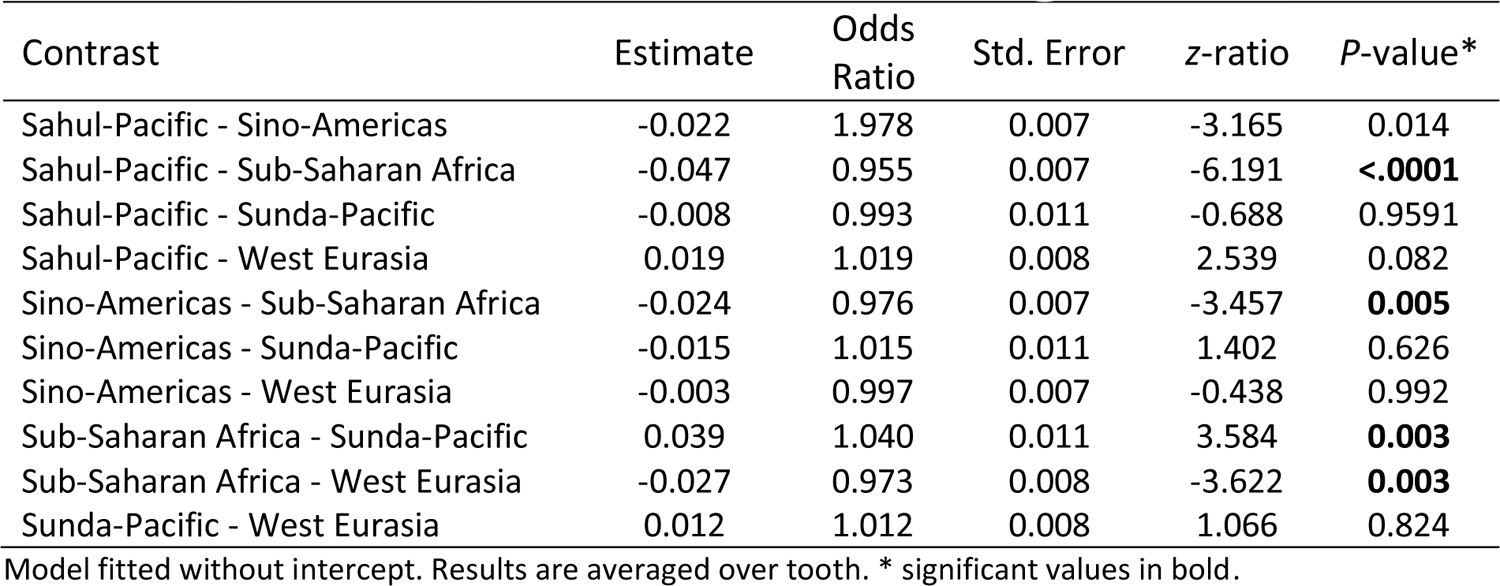
Tukey pair-wise comparisons of canal to root number by geographical region

